# Neural extracellular matrix regulates visual sensory motor integration

**DOI:** 10.1101/2023.04.17.537074

**Authors:** Jacqueline Reinhard, Cornelius Mueller-Buehl, Susanne Wiemann, Lars Roll, Veronika Luft, Hamed Shabani, Daniel L. Rathbun, Lin Gan, Chao-Chung Kuo, Julia Franzen, Stephanie C. Joachim, Andreas Faissner

**Affiliations:** Department of Cell Morphology and Molecular Neurobiology, Faculty of Biology and Biotechnology, Ruhr University Bochum, 44780 Bochum, Germany; Institute for Ophthalmic Research, Centre for Ophthalmology, Eberhard-Karls-University Tuebingen, 72076 Tuebingen, Germany; Interdisciplinary Centre for Clinical Research Aachen, RWTH Aachen University, 52074 Aachen, Germany; Experimental Eye Research Institute, University Eye Hospital, Ruhr University Bochum, 44892 Bochum, Germany

**Keywords:** brevican, direction-selective cholinergic starburst amacrine cells, direction-selective ganglion cells, extracellular matrix, neurocan, retina, synapse, tenascin-C, tenascin-R, visual motion processing

## Abstract

Visual processing depends on sensitive and balanced synaptic neurotransmission. Extracellular matrix proteins in the environment of cells are key modulators in synaptogenesis and synaptic plasticity. In the present study, we provide evidence that the combined loss of the four extracellular matrix components brevican, neurocan, tenascin-C and tenascin-R in quadruple knockout mice leads to severe retinal dysfunction and diminished visual motion processing *in vivo*. Remarkably, impaired visual motion processing was accompanied by a developmental loss of cholinergic direction-selective starburst amacrine cells. Additionally, we noted imbalance of inhibitory and excitatory synaptic signaling in the quadruple knockout retina. Collectively, the study offers novel insights into the functional importance of four key extracellular matrix proteins for retinal function, visual motion processing and synaptic integrity.

**Graphical Abstract:** 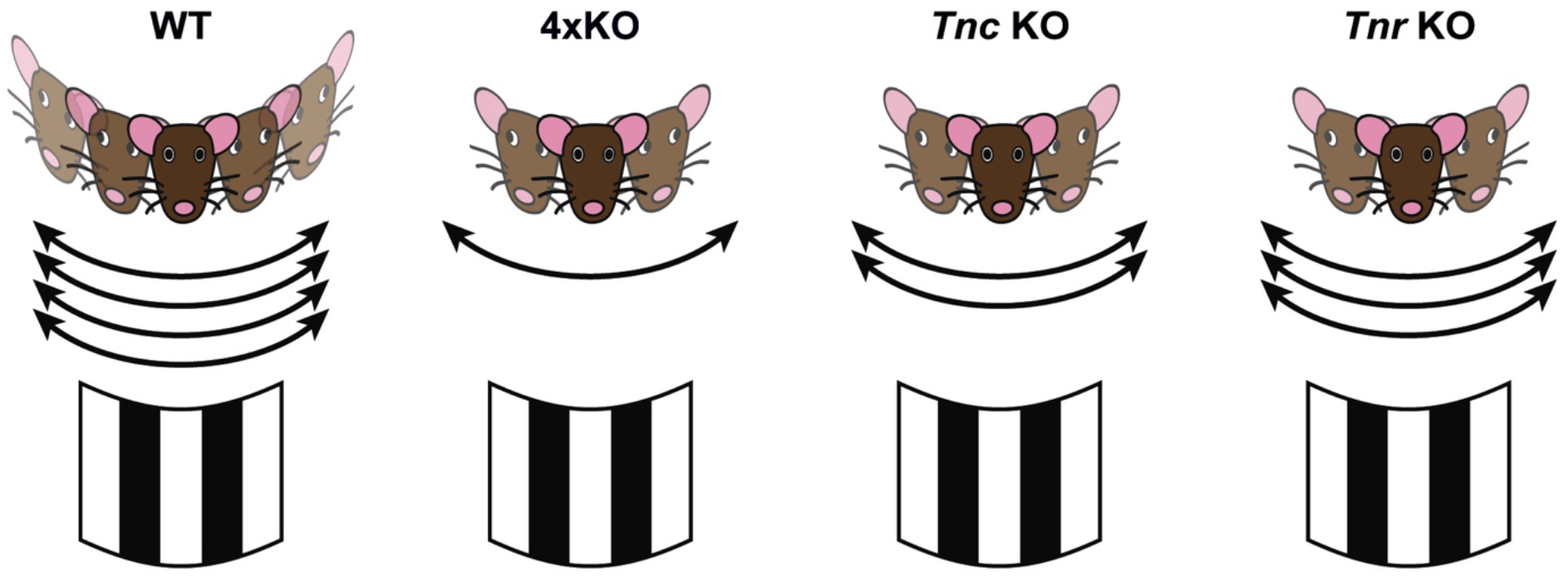

**In brief:** In their study, Reinhard et al. show that the combined loss of the extracellular matrix components brevican, neurocan, tenascin-C and tenascin-R in quadruple knockout mice leads to retinal dysfunction, diminished visual motion processing, developmental loss of cholinergic direction-selective starburst amacrine cells and imbalance of inhibitory and excitatory synaptic integrity.

**Highlights:**

- Combined loss of the four extracellular matrix molecules brevican, neurocan, tenascin-C and tenascin-R causes retinal dysfunction
- Impaired visual motion processing in quadruple, tenascin-C and tenascin-R knockout mice
- Loss of cholinergic direction-selective starburst amacrine cells in the quadruple knockout retina
- The matrisome influences inhibitory and excitatory synaptic balance

## INTRODUCTION

Growing evidence indicates that the extracellular matrix (ECM) matrisome and interacting molecules are key players in synaptic maturation and plasticity [1, 2]. The ECM is a highly organized and interactive meshwork composed of complexes of hyaluronan and proteoglycans linked by oligomeric glycoproteins [3]. Particularly, ECM molecules of the tenascin glycoprotein family have become a research focus as important synaptic modulators [4, 5]. For example, tenascin-C (Tnc) modulates the activity of L-type Ca^2+^ channels [6] and is expressed in specific patterns in the developing central nervous system (CNS). Thereafter, Tnc is downregulated but persists in regions of neuronal plasticity, such as the hippocampus [7]. The tenascin family member tenascin-R (Tnr) is an important component of perineuronal nets (PNNs) [8]. Thus, the loss of Tnr leads to an abnormal PNN formation and an altered distribution of PNN-associated ECM molecules in *Tnr* knockout (KO) mice [9]. Important binding partners of tenascins are chondroitin sulfate proteoglycans (CSPGs), which consist of a core protein with covalently attached glycosaminoglycan chains [10, 11]. The CNS-specific CSPGs neurocan (Ncan) and brevican (Bcan) inhibit neurite outgrowth [12, 13] and display developmentally regulated expression in the rodent CNS [14]. Both tenascins as well as CSPGs have been identified as structural components of PNNs [15, 16]. Furthermore, the elimination of either *Bcan*, *Tnc* or *Tnr* in gene KO approaches resulted in the modification of long-term potentiation (LTP) in the CA1 region [6, 17–19].

The integration of visual motion processes relies on a precise and finely tuned synaptic neurotransmission. In the retina, starburst amacrine cells (SACs) play a pivotal role in direction-selectivity and the detection of directional motion [20, 21]. SACs associate with the retinal ganglion cell layer (GCL) and inner nuclear layer (INL) and are characterized by the expression of choline acetyltransferase (ChAT). By releasing γ-aminobutyric acid (GABA), SACs modulate synaptic input on direction-selective RGCs (DSGCs) [22]. Notably, GABA_A_ receptors containing the α 2 subunit have been found to be critical for direction-selective inhibition [23].

To examine the role of the neural ECM in visual motion processing, we studied mice lacking the ECM components Bcan, Ncan, Tnc and Tnr. These quadruple KO mice are viable and fertile [24]. Previous investigations revealed that hippocampal neurons from quadruple KO mice exhibit diminished PNNs, altered expression of synaptic proteins and lower miniature excitatory and inhibitory postsynaptic current (mEPSC/mIPSC) frequencies [25]. A reduced short-term depression and changed frequency dependence was observed in the quadruple KO hippocampus *in vivo* [26], paralleled by a dramatic change in the ratio of excitatory and inhibitory synapses in cultured hippocampal neurons [27]. Moreover, these ECM proteins are important regulators of the interplay between PNNs, synaptic integrity, inhibitory interneurons and the transcription factor Otx2 (orthodenticle homeobox 2) in the visual cortex [28]. These observations indicate that the neural ECM preserves the equilibrium of neuronal networks by stabilizing inhibitory synapses [29]. However, it is currently poorly understood if and to which extent the quadruple KO results in an excitatory-inhibitory synaptic imbalance *in vivo*. It is also not known which functional and cellular consequences this entails in neural networks. In this perspective, we focused on retinal function in quadruple KO mice. So far, the consequences of an ECM KO for visual processing are unknown [30, 31].

Herein, we demonstrate that retinal physiology is impaired in the quadruple KO mice, suggesting severe retinal dysfunction. Most interestingly, altered visual motion processing was evident in the quadruple KO as well as in *Tnc* or *Tnr* single KO mice. This could be traced to a loss of cholinergic direction-selective SACs, an impaired balance of inhibitory GABAergic and excitatory glutamatergic synapses and significant changes in the transcriptome of quadruple KO retinae. Collectively, our study provides first evidence that the ECM influences visual sensory motor behavior already at the retina level, without intervention of disease and/or experimental manipulation.

## RESULTS

### Differential expression of Bcan, Ncan, Tnc and Tnr in the postnatal and adult mouse retina

Before analyzing the functional importance of Bcan, Ncan, Tnc and Tnr, we wanted to elucidate their detailed expression patterns during mouse retinal development via *in-situ* hybridization and immunohistochemistry (Figure 1; Figure S1; Figure S2). The expression of Bcan in the developing mouse retina has not been analyzed so far. *Bcan* mRNA expression was hardly detectable at postnatal day (P) 0 and P4 but increased throughout ongoing retinal development (Figure S1A-È).

**Figure 1.**
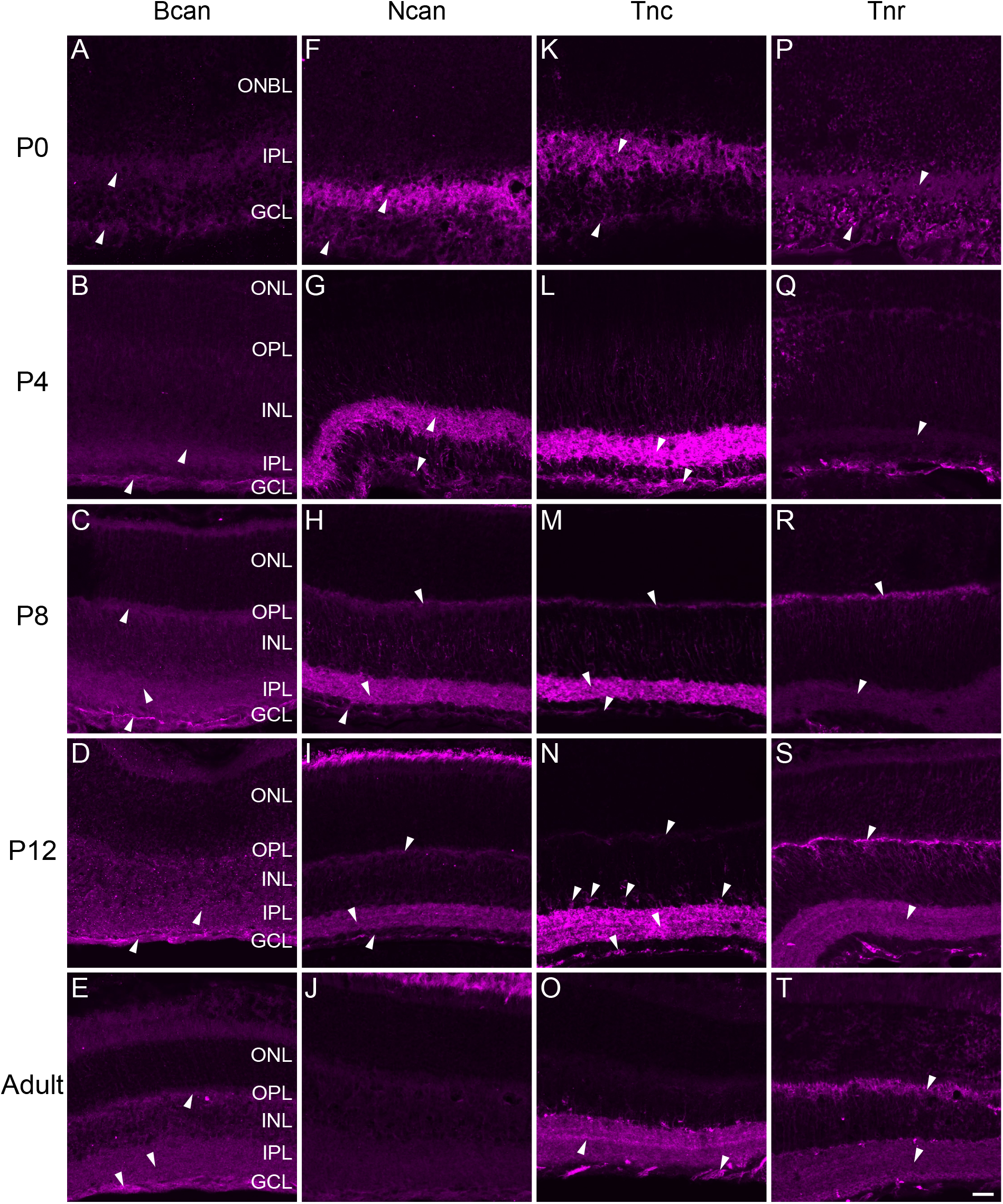
Immunohistochemical staining of Bcan, Ncan, Tnc and Tnr in the postnatal and adult mouse retina. Differential staining pattern of Bcan (A-E), Ncan (F-J), Tnc (K-O) and Tnr (P-T) in the postnatal (P0, P4, P8 and P12) and adult retina. White arrows mark prominent signals. Scale bar: 20 µm. Bcan = brevican; GCL = ganglion cell layer; INL = inner nuclear layer; IPL = inner plexiform layer; Ncan = neurocan; ONBL = outer neuroblastic layer; ONL = outer nuclear layer; OPL = outer plexiform layer; P = postnatal; Tnc = tenascin-C; Tnr = tenascin-R.

Also, on protein level, we observed weak immunoreactivity of Bcan in the inner plexiform layer (IPL) and the GCL around P0 and P4 (Figure 1A, B). From P8 onwards, we found signals in the outer plexiform layer (OPL) and the immunoreactivity in the GCL increased (Figure 1C-E).

The protein pattern of Ncan was described previously in the developing rat retina [32]. When we analyzed the mRNA expression of *Ncan* in the P0 mouse retina, we observed faint signals throughout all retinal layers (Figure S1F-J′). At P4, these signals became more pronounced, especially in the inner retina. However, *Ncan* mRNA expression was less distinct in the adult retina. The immunohistochemical analyses revealed Ncan protein from P0 onwards in the IPL and GCL (Figure 1 F-J). At P8, Ncan was also observed in the OPL. At adulthood, no specific Ncan immunoreactivity was observed. Remarkably, Ncan signals vanished in retinal layers.

Developmental Tnc expression has been previously reported in the chick retina [33]. Here, *Tnc* mRNA was detected in amacrine cells (ACs), displaced ACs and horizontal cells (HCs). In our study, we revealed a similar *Tnc* expression in the developing mouse retina. *Tnc* mRNA expression was detected in single cells of the outer neuroblastic layer (ONBL) at P0 (Figure S1K, K′). From P4 until P12, Tnc signals were found at the basal side of the INL and in the GCL, where ACs are located (Figure S1L-N′). This expression pattern was also noted in the adult retina (Figure S1O, Ò). Additionally, we observed signals at the apical side of the INL, which seemed to be HCs. On protein level, Tnc was localized in the IPL and GCL from P4 onwards (Figure 1K-O). At P8, additional signals were observed in the OPL. In the adult retina, Tnc immunoreactivity decreased slightly.

Tnr expression was previously described in HCs. In the mouse retina, it seems to increase until the third postnatal week and then remains at a constant level [34]. Our results suggest that some cells in the ONBL already expressed *Tnr* mRNA at birth (Figure S1P-P′). Upon P4, *Tnr* mRNA was expressed by cells at the apical side of the INL, which showed HC morphology (Figure S1Q-T′). Signals were also observed at the basal side of the INL and in the GCL. On protein level, only faint Tnr signals were detected at P0 and P4 (Figure 1P, Q). After P8, Tnr immunoreactivity highly increased in the OPL as well as in the IPL and this expression pattern was maintained until the adult stage (Figure 1R-T).

Taken together, our study showed that the expression of Bcan, Ncan, Tnc and Tnr is developmentally regulated in the mouse retina. Most interestingly, on protein level, these ECM constituents showed a prominent expression in the plexiform layers and nerve fiber layer beneath the RGCs, which points to their potential functional relevance at synaptic sites.

### Physiological intraocular pressure but severe reduction of a-/b-wave amplitudes in quadruple KO mice

To explore possible physiological and functional alterations in quadruple KO mice, we measured the intraocular pressure and analyzed the retinal function. Tonometry analyses revealed a comparable and physiological intraocular pressure in wildtype (WT) and quadruple KO mice (quadruple KO: 11.39 ± 0.60 mmHg; WT: 11.94 ± 0.52 mmHg; p = 0.49; Figure 2A).

**Figure 2.**
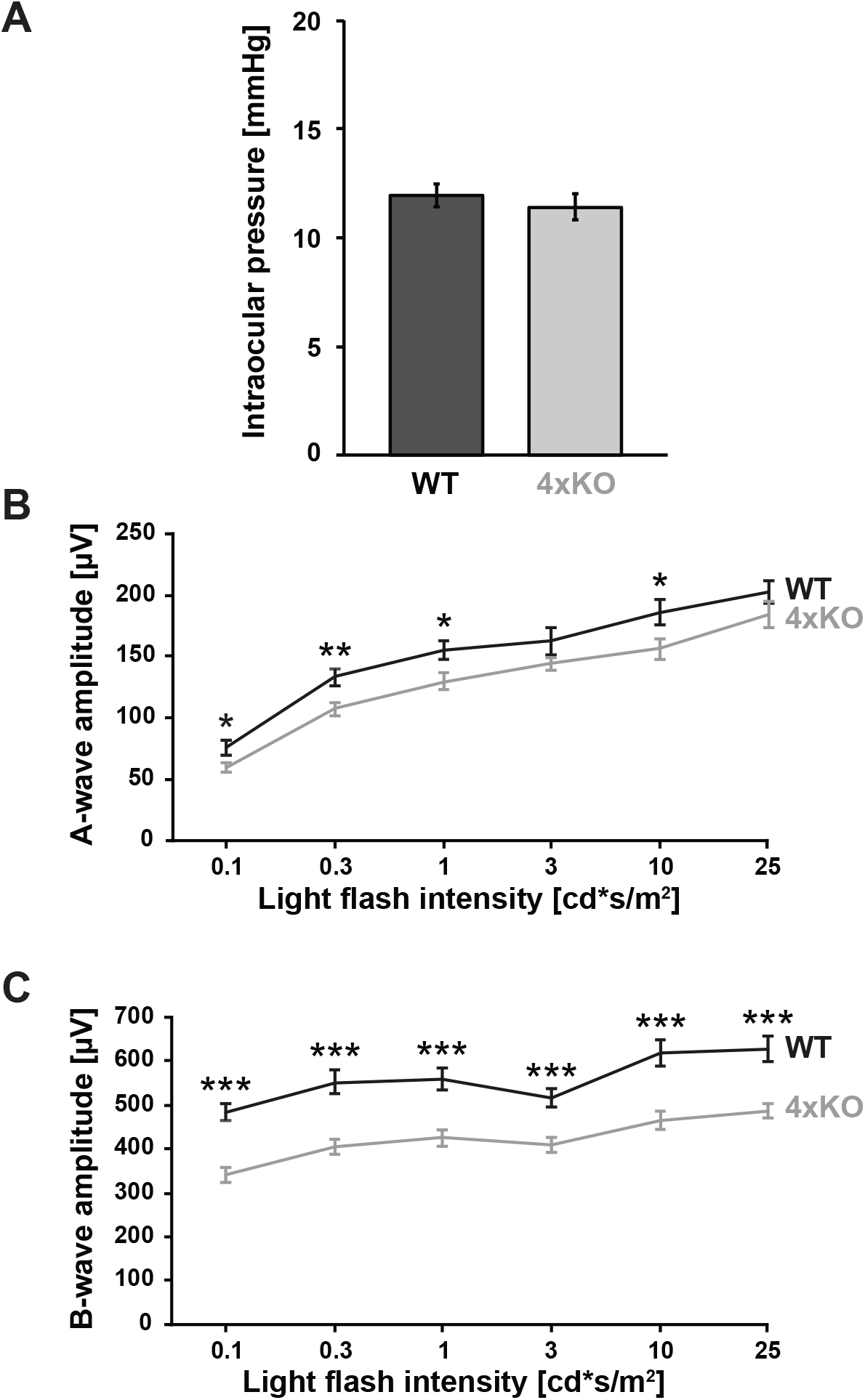
Physiological intraocular pressure but severe reduction of a-/b-wave amplitudes in quadruple KO mice. (A) Intraocular pressure of both eyes was measured in quadruple KO and WT mice. Both, quadruple KO and WT mice, exhibited a comparable, normal physiological intraocular pressure. (B, C) ERG recordings revealed significantly reduced a-wave amplitudes in quadruple KO compared to WT animals. Remarkably, b-wave amplitudes were significantly decreased. * = p < 0.05, ** = p < 0.01, *** = p < 0.001; N = 8. 4xKO = quadruple KO, WT = wildtype. Data are shown as mean ± SEM.

Functional analyses were performed by scotopic electroretinogram (ERG) recordings (Figure 2B-C, Table S1). Here, the a-wave represents electrical responses of the rod photoreceptors, while the b-wave shows electrical responses of BCs to the light flash. ERG recordings showed that quadruple KO mice had significantly reduced a-wave amplitudes (p < 0.05 at 0.1, 0.3, 1 and 10 cd*s/m^2^; Figure 2B; Table S1). Most notably, b-wave amplitudes were reduced (p < 0.001; 0.1 to 25 cd* s/m^2^; Figure 2C; Table S1). These data indicated severe retinal functional deficits in quadruple KO mice, which seemed to particularly affect the inner retina.

### Impairment of optomotor responses in quadruple KO mice as well as *Tnc* and *Tnr* single KO mice

Based on our findings of an impaired retinal function in quadruple KO mice, we implemented further measurements of visual motion processing. Therefore, we measured the optomotor response (OMR), a basic mechanism to stabilize images on the retina in a moving environment [35]. We analyzed the OMR in quadruple KO as well as in *Tnc* and *Tnr* single KO mice (Figure 3A-D). As measure for the OMR, we counted the number of saccades, a fast-returning movement of the head after following the moving stripes to stabilize the image on the retina. Quadruple KO mice showed a significantly reduced number of saccades/minute at slow (20^°^/second) as well as at fast velocities (50^°^/second) and in both directions (rotation clockwise and counterclockwise) compared to WT mice (p < 0.001).

**Figure 3.**
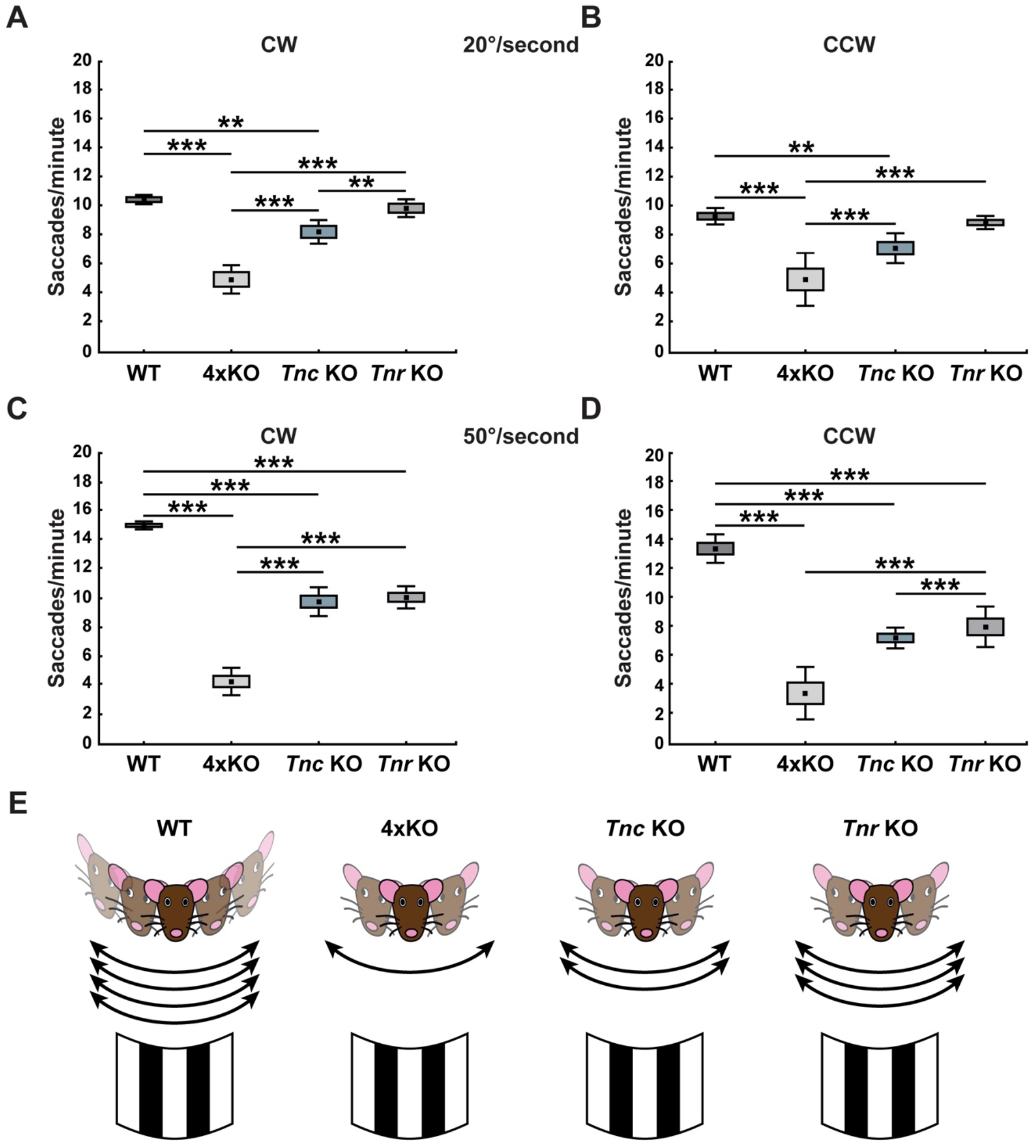
Impairment of optomotor responses in quadruple KO mice as well as *Tnc* and *Tnr* single KO mice. (A-D) quadruple KO mice showed a significantly reduced number of saccades/minute at slow (20^°^/second; A and B) as well as fast velocities (50^°^/second; C, D) and in both directions (rotation clockwise (CW), A, C; and rotation counterclockwise, CCW, B, D) compared to WT mice. Additionally, quadruple KO mice displayed a lower saccade number in comparison to *Tnc* KO and *Tnr* KO animals. Compared to the WT group, as shown for both directions and velocities, *Tnc* KO mice exhibited less saccades/minute. *Tnr* KO animals displayed a comparable number of saccades/minute at slow velocities in both directions compared to control mice. However, a significant reduction was observed at fast velocities. Only a slightly reduced or comparable saccade number was found in *Tnc* KO when compared to *Tnr* KO mice. (E) As schematically shown, these data demonstrate a defective direction-selectivity in the investigated ECM mutants, which was most severe in quadruple KO mice. ** = p < 0.01, *** = p < 0.001; N = 6. 4xKO = quadruple KO, CW = clockwise, CCW = counterclockwise, *Tnc* KO = *Tnc* knockout, *Tnr* KO = *Tnr* knockout, WT = wildtype. Data are shown as median ± quartile + minimum/maximum.

Quadruple KO mice also displayed a lower saccade number in comparison to *Tnc* (p < 0.01) and *Tnr* (p < 0.001) KO animals. Compared to the WT group, as shown for both directions, *Tnc* KO mice exhibited less saccades at 20^°^/second (p < 0.01) and 50^°^/second (p < 0.001). *Tnr* KO animals showed a comparable number of saccades at slow velocities in both directions as WT mice (p > 0.05). However, a significant reduction was observed at fast velocities (p < 0.001). Only a slightly reduced (p < 0.05; 20^°^/second, clockwise) or comparable saccade number (p > 0.05; 20^°^/second, CCW; 50^°^/second, clockwise and counterclockwise) was found in *Tnc* KO mice when compared to *Tnr* KO animals.

In summary, our data indicate a defective visual processing in the investigated ECM mutants, which was most severe in quadruple KO mice. Interestingly, compared to quadruple KO mice, *Tnc* single KO mice showed less severe impairment in the optomotor response behavior, but significant limitations compared to *Tnr* KO mice. These results indicate that the four ECM molecules are critical for visual motion processing.

### Loss of cholinergic direction-selective ON-SACs in the quadruple KO retina

Because of the reduced number of saccades in quadruple KO mice, we next analyzed whether these defects are associated with deficits on the cellular level. Cholinergic direction-selective SACs play a crucial role in mediating retinal direction-selectivity. Thus, destruction of SACs leads to a loss of directionality in RGCs as well as a loss of optokinetic responses, indicating their importance for stabilizing image motion [36, 37]. Additionally, selective ablation of SACs leads to a loss of OMRs [38].

To analyze the number of SACs in quadruple KO and WT retinae, flat mounts were immunostained with anti-ChAT antibodies. Thus, cell bodies of ON- and OFF-cholinergic SACs were specifically stained in the GCL and INL, respectively (Figure 4A-B′) [39]. Quantification revealed a significantly lower number of cholinergic SACs in the GCL of quadruple KO compared to WT retinae (WT: 653.5 ± 134.6 cells/mm^2^; quadruple KO: 474.9 ± 86.0 cells/mm^2^, p = 0.01, Figure 4C). In contrast, no change in the number of ChAT-positive cells could be observed in the INL (WT: 729.8 ± 99.7 cells/mm^2^; quadruple KO: 806.1 ± 111.0 cells/mm^2^, p = 0.2, Figure 4D). These results indicated a specific loss of cholinergic ON-SACs in the quadruple KO retina.

**Figure 4.**
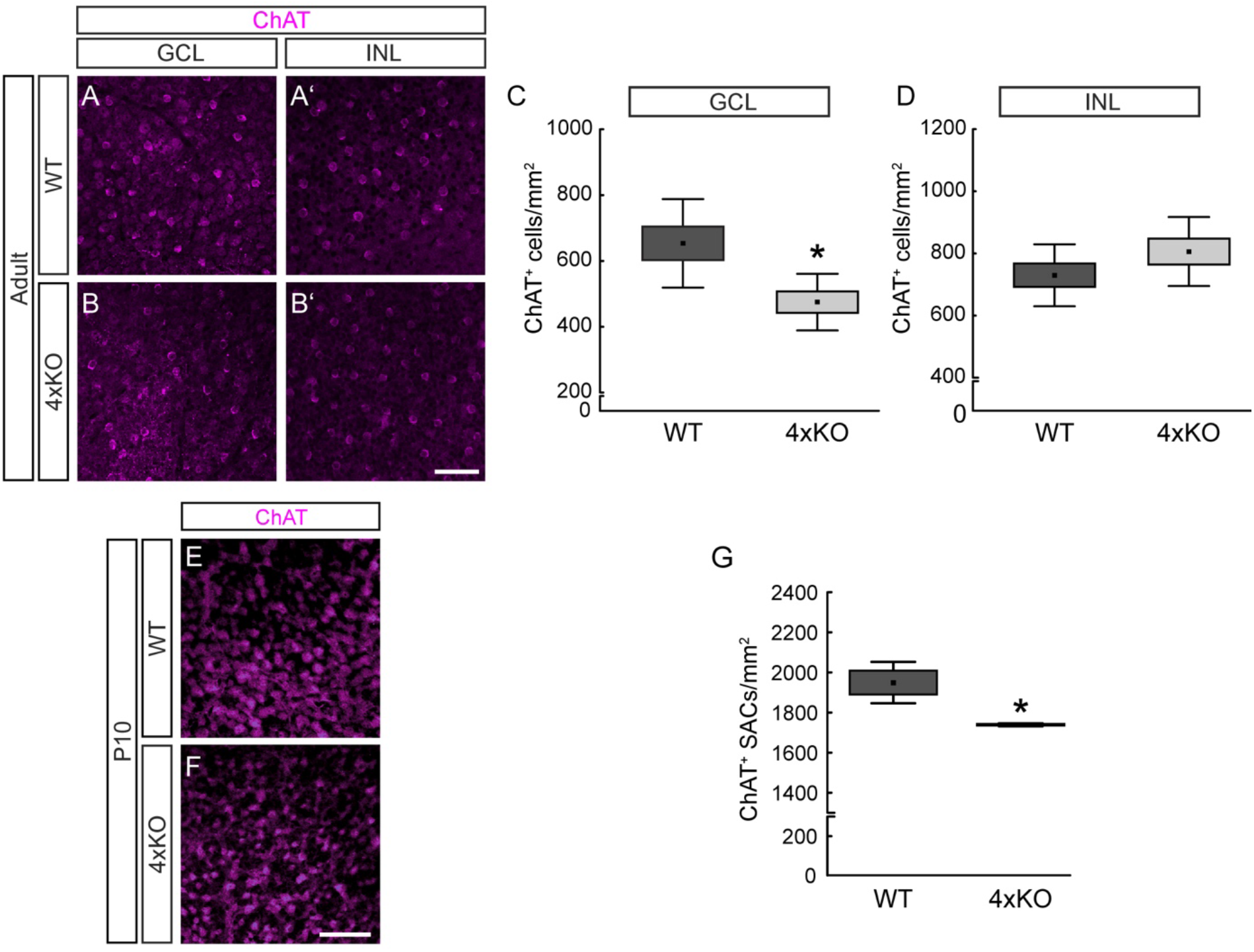
Loss of cholinergic direction-selective ON-SACs in the quadruple KO retina. (A-B′) Representative photographs of ChAT-positive SACs (magenta) in the GCL (= ON-SACs; A, B) and INL (= OFF-SACs; À, B′) of WT and quadruple KO flat mount retinae. (C, D) The number of ON-/OFF-ChAT-positive cells was determined in the GCL (C) and INL (D). A significant loss in the number of ON-ChAT-positive cells was found in quadruple KO compared to control mice (C, p = 0.01). No alterations of OFF-ChAT-positive cells could be noted (D, p = 0.2). N = 7. (E, F) Exemplary photographs of ChAT-positive SACs (magenta) in P10 flat mount quadruple KO and WT retinae. (G) Quantification revealed a significantly reduced number of ChAT-positive SACs in postnatal quadruple KO retinae (p < 0.05). N = 3. * = p < 0.05; ChAT = choline acetyltransferase, GCL = ganglion cell layer, INL = inner nuclear layer; ON-SAC = ON-starburst amacrine cell; 4xKO = quadruple knockout, WT = wildtype. Data are shown as mean ± SEM and SD. Scale bars = 50 μm.

In the mouse retina, acetylcholine (ACh) is required for the propagation of spontaneous activity after birth to P10 [40]. Around eye opening at P12, cholinergic neurons show a mature-like phenotype [41]. To investigate whether the loss of ChAT-positive SACs in quadruple KO retinae was already evident in early development, we stained retinal flat mounts at P10 (Figure 4E-F). At this point in time, migrating SACs cannot be clearly assigned to the INL or GCL. Therefore, we counted the total number of SACs in both layers of the retina. We noted a significant reduction in premature total ChAT-positive SACs from quadruple KO retinae (WT: 1949.5 ± 59.5 cells/mm^2^; quadruple KO: 1739.0 ± 4.0 cells/mm^2^, p = 0.02, Figure 4G). Based on these results, we conclude that the reduced number of SACs in quadruple KO already originates during early retinal development.

### Comparable response of DSGCs in quadruple KO and WT mice

Direction-selectivity of DSGCs results from patterned excitatory and inhibitory inputs during motion stimuli. A critical factor for DSGC direction-selective processing is their inhibition *via* SACs. Given the reduction in the number of SACs in the GCL of quadruple KO retinae, it was of great interest to analyze the electrical DSGC response to a moving object. Thus, to investigate possible changes in DSGC responses, MEA recordings, with a light bar moving in different directions as stimulus, were carried out in quadruple KO and WT retinae (Figure 5A-F, Figure S3). The polar plot (Figure 5A) and peristimulus time histogram (PSTH; Figure 5 B) visualize the exemplary response of an individual DSGC. The cell mainly responded to a movement of the light bar at an angle of 45° and 180° (Figure 5B-D). In the other directions, the cell hardly reacted. Overall, we noted a tendency towards lower DSGC responses in quadruple KO compared to WT mice (Figure 5E-F). However, statistical analyses revealed comparable DSGC responses in both genotypes (p > 0.05).

**Figure 5.**
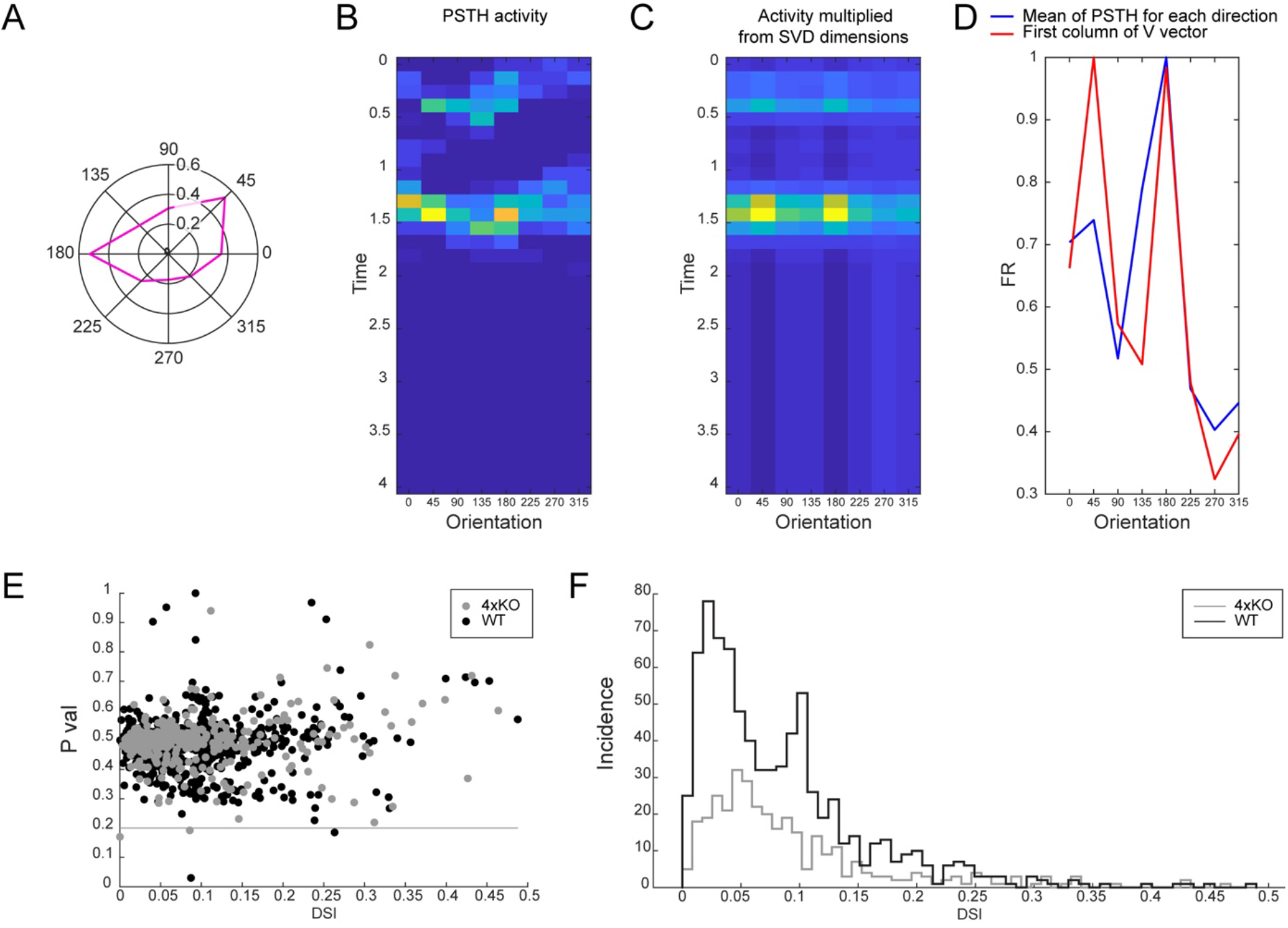
No significant change in the response of DSGCs. DSGC responses were recorded in quadruple KO and WT retinae by MEA analyses. (A) The polar plot and (B) peristimulus time histogram (PSTH) show the exemplary response of an individual DSGC (magenta in A). The cell mainly responded to movement of the light bar at an angle of 45^°^ and 180° (yellow/orange in B). (C) Projected data using the first column of V, drawn from SVD decomposition of the PSTH response matrix shown in B. (D) Red: average of PSTH in B, blue: first column of V. (E) The dot diagram displays the p-value and the direction-selectivity index of WT DSGCs (black dots) and quadruple KO DSGCs (grey dots). (F) The diagram depicts the cumulative responses (incidence) of WT DSGCs (black line) and quadruple KO DSGCs (grey line). Although we observed a trend towards lower DSGC responses in quadruple KO compared to WT mice, no significant differences were found between both genotypes. N = 10. DSGC = direction-selective retinal ganglion cell; DSI = direction-selectivity index, FR = firing rate, 4xKO = quadruple knockout, WT = wildtype.

### Imbalance of inhibitory and excitatory synaptic integrity in the quadruple KO retina

SACs generate direction-selective output of GABA to provide critical inhibition of receptive field properties of DSGCs. As previously shown, GABA_A_ receptors containing the α2 subunit are critical for direction-selective inhibition [23]. Interestingly, our analyses revealed that quadruple KO mice exhibited a significantly reduced number of inhibitory GABA_A_ receptor-positive cells in the GCL (WT: 92.5 ± 2.4 cells/mm; quadruple KO: 84.5 ± 1.6 cells/mm, p < 0.02, Figure 6A-C).

**Figure 6.**
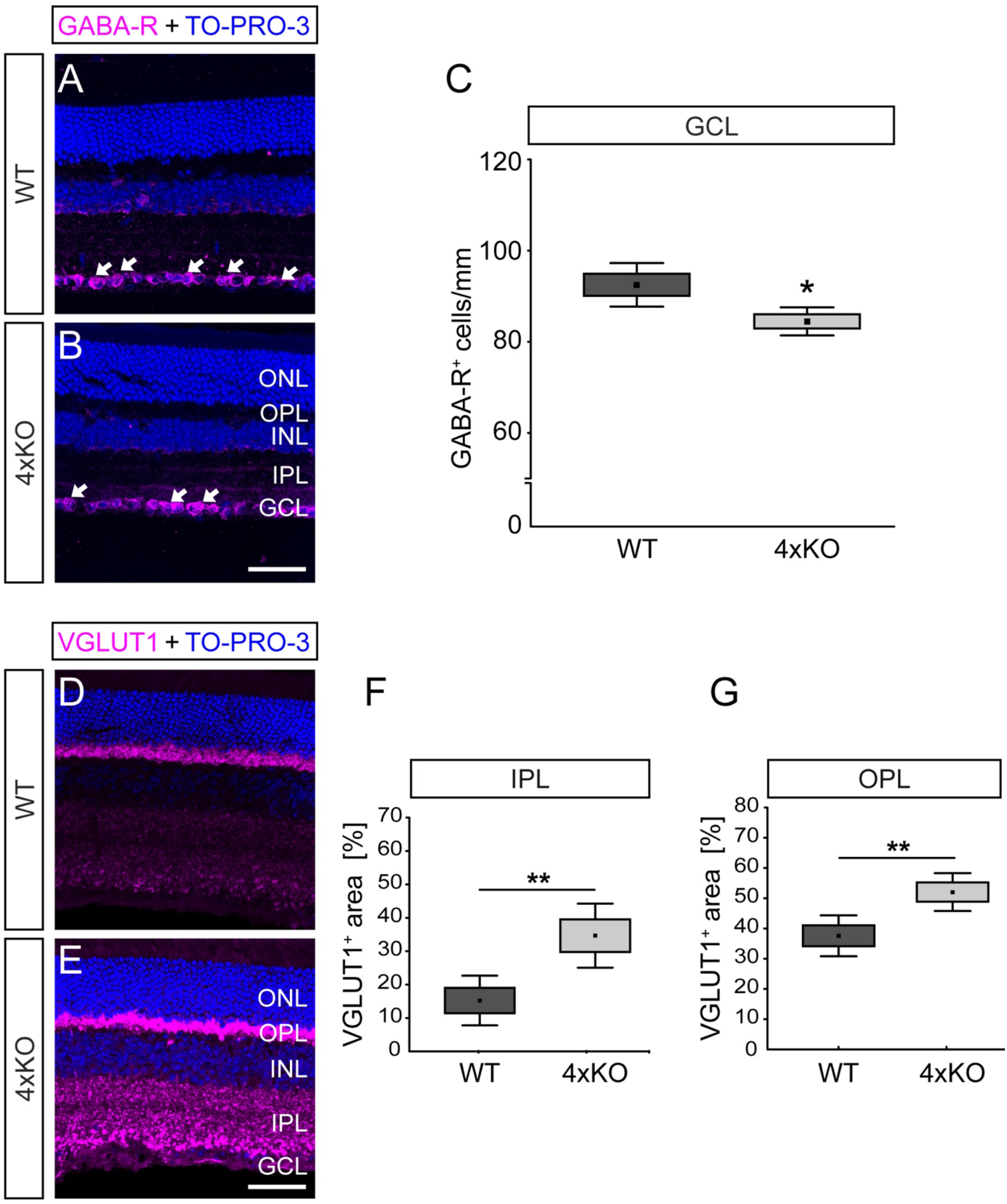
Imbalance of inhibitory and excitatory synaptic integrity in the quadruple KO retina. (A, B) Representative images of retinal sections of WT and quadruple KO mice, which were stained using anti-GABA-R (GABA_A_ receptor α2, magenta). Cell nuclei were stained with TO-PRO-3 (blue). (C) Quantification revealed a significantly reduced number of GABA-R-positive cells in the GCL of quadruple KO mice (p < 0.05). * = p < 0.05; N = 8. (D, E) Exemplary images of anti-VGLUT1 (magenta) stained retinal sections of WT and quadruple KO mice. (F, G) quadruple KO mice showed a significantly enhanced excitatory VGLUT1 staining in the plexiform layers (IPL and OPL). Cell nuclei were stained with TO-PRO-3 (blue). ** = p < 0.01; N = 8. GABA-R = γ-aminobutyric acid receptor A, GCL = ganglion cell layer, INL = inner nuclear layer, 4xKO = quadruple knockout, VGLUT1 = vesicular glutamate transporter 1, WT = wildtype. Data are shown as mean ± SEM and SD. Scale bars = 50 μm.

Inhibition is essential to counterbalance excitatory neurotransmission. Excitatory glutamatergic neurotransmission depends on vesicular glutamate transporters (VGLUTs), which segregate glutamate into synaptic vesicles. As previously shown, VGLUT1 is required for photoreceptor signaling to second- and third-order neurons, but not for intrinsic visual functions [42]. The immunoreactivity pattern of VGLUT1 is restricted to photoreceptor and BC terminals, the principle glutamatergic neurons in the retina [43, 44].

To comparatively assess VGLUT1 staining in WT and quadruple KO retinae, retinal cross-sections were stained with anti-VGLUT1 antibodies. Representative images show a specific synapse-associated VGLUT1 staining in both plexiform layers (Figure 6D-E). However, as validated by quantification, quadruple KO mice showed a significantly larger VGLUT1 staining area in the IPL compared to WT mice (WT: 13.9 ± 8.6% VGLUT1 positive area; quadruple KO: 37.2 ± 10.5% VGLUT1 positive area, p < 0.001; Figure 6F). In addition, we found a significantly increased VGLUT1 positive area in the OPL of quadruple KO in comparison to WT mice (WT: 37.6 ± 9.8% VGLUT1 positive area vs. quadruple KO: 52.0 ± 9.0% VGLUT1 positive area, p = 0.008; Figure 6G) pointing to an increased excitatory signaling. Taken together, our findings suggest quadruple KO mice exhibit an altered balance of retinal excitatory/inhibitory neurotransmission *in vivo*.

### Global gene expression changes in the quadruple KO retina

To obtain a more detailed picture on global gene expression changes in the adult quadruple KO retina, we performed NGS analyses. Subsequent bioinformatic analyses identified a total of 263 differentially expressed genes (adjusted p < 0.05) of which 142 genes were downregulated and 121 genes were upregulated in the quadruple KO compared to the WT retina (Table S2). The heatmap and volcano plot illustrate the identified significant differentially expressed genes (Figure S4A, B). Additionally, heatmaps show row z-score scaled expression values of regulated genes (adjusted p < 0.05), which belong to enriched Gene Ontologies (GOs) in three function groups of interest: *Extracellular matrix* (13 genes), *Visual function* (44 genes) and *Development* (60 genes) as well as the selected GOs in function group S*ynapse* (17 genes) (Figure 7A-D).

**Figure 7.**
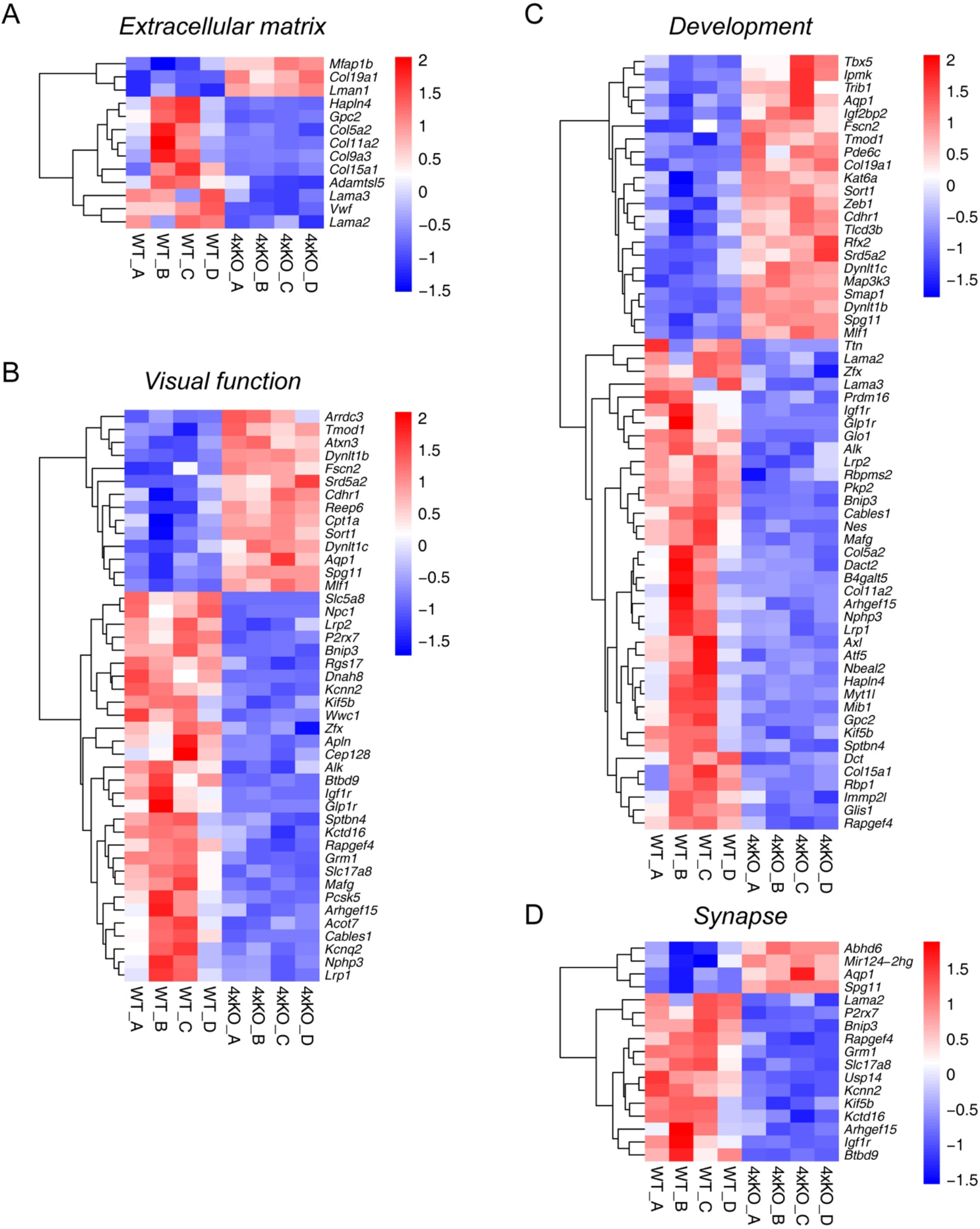
NGS revealed global gene expression changes in the quadruple KO retina. (A-D) NGS revealed a different gene expression pattern in the quadruple KO compared to the WT mouse (N = 4). The heatmaps include significantly regulated genes that belong to the three enriched GOs in function groups of interest (A) *ECM*, (B) *Visualfunction* and (C) *Development* as well as for the selected GOs in function group of interest (D) *Synapse*. Heatmaps present the normalized expression (log2 scale) of significantly regulated genes (adjusted p-value < 0.05) annotated with GO terms for the four groups of interest. The expression change of these genes is indicated by the color shift from blue to red. Thus, downregulated genes are illustrated in blue, while upregulated genes are depicted in red. KO = quadruple knockout, WT = wildtype.

Interestingly, we detected a dysregulation of various ECM and ECM-associated genes in the quadruple KO retina. A significant downregulation was noted for *Hapln4 (hyaluronan and proteoglycan link protein 4)*, the ECM-receptor *Gpc2 (glypican 2)*, the *α collagens Col5a*, *Col9a3, Col11a2* and *Col15a1*, *Adamtsl5* (*a disintegrin-like and metalloprotease domain containing thrombospondin type 1 motif-like 5*) as well as the *α laminins Lama2* and *Lama3*. In contrast, *Mfap1b (microfibrillar-associated protein 1B)*, *Col19a1* and *Lman 1 (lectin, mannose-binding 1)* were significantly upregulated in the quadruple KO retina. Besides, we noted a dysregulation (raw p-value lower than 0.05) of ECM/ECM-related genes such as *α collagens Col6a1*, *Col1a1*, *Col2a1*, *laminins Lama1, Lamb2*, the ECM-modifying gene *Timp2 (tissue inhibitor of metalloproteinase 2)* and the ECM-receptors *Itgb4* (*integrin β4)*, *Gpc1 (glypican 1)* as well as *CD44* (cluster of differentiation 44; data not shown).

Additionally, GO term analyses revealed an enrichment of the function group *Visual function*, which includes the upregulation of *Arrdc3 (arrestin domain containing protein 3*) as well as the AC and Müller glia cell-expressed water channel *Aqp1* (*aquaporin 1*) [45]. In contrast, a downregulation was observed for genes such as *Kcnq2 (potassium voltage-gated channel, subfamily Q, member 2)*, *Lrp1* and *Lrp2* (*low density lipoprotein receptor-related protein 1 and 2*) as well as *Glp1r (glucagon-like peptide 1 receptor).* Among others, enrichment analyses of the function group *Development* showed a downregulation of the RGC-specific gene *Rbpms2* (*RNA binding protein with multiple splicing 2*) [46].

Based on previous knowledge and data of the present work, we were also interested in the dysregulation of genes annotated to the selected function group *Synapse.* Most interestingly, we observed a dysregulation of several synaptic proteins, which included a prominent upregulation of the gene *Abhd6* (*abhydrolase domain containing 6*), the long non-coding RNA *Mir124-2hg*, *Aqp1* (see also GO groups *Visual function* and *Development*) as well as *Spg11* (*spatacsin vesicle trafficking associated*).

A significant downregulation was found, e.g., for the synaptic/synapse-associated genes Lama2, P2rx7 (purinergic receptor P2X, ligand-gated ion channel 7), Bnip3 (BCL2/adenovirus E1B interacting protein 3), Rapgef4 (Rap guanine nucleotide exchange factor (GEF) 4), Grm1 (glutamate receptor, metabotropic 1), Slc17a8 (sodium-dependent inorganic phosphate cotransporter, member 8; also known as the vesicular glutamate transporter 3/Vglut3), Kcnn2 (potassium intermediate/small conductance calcium-activated channel, subfamily B, member2), Kctd16 (potassium channel tetramerization domain containing 16), Igf1r (insulin-like growth factor I receptor) and Btbd9 (BTB (POZ) domain containing 9).

Collectively, we noted an extensive ECM remodeling and synaptic imbalance in quadruple KO mice, which might directly contribute to the observed functional and optomotor deficits.

## DISCUSSION

In the 1960′s, direction-selective nerve cells in the retina of vertebrates, which only become activated when a light stimulus completes a specific movement, were first described [47]. These cells, known today as SACs, play a critical role in inhibitory GABAergic and excitatory cholinergic modulation of DSGC responses, contributing to directional selectivity and the recognition of directional motion [48]. In our study, we report on the impairment of visual motion processing and loss of cholinergic direction-selective SACs in quadruple KO mice lacking the four ECM proteins Bcan, Ncan, Tnc and Tnr. A reduced number of SACs in the quadruple KO at P10 suggests that the loss of the ECM constituents already impacts early development of SACs. In this regard, the specification, maturation, migration and/or survival of SACs might be affected. Accordingly, tenascins and CSPGs have been shown to provide a crucial modulatory environment for those cellular functions [49–52]. We cannot exclude that also other cell types are affected by the loss of the four ECM molecules. However, particularly the loss of Tnc expression in SACs might be directly related to a cell intrinsic mechanism. Our analyses in the WT retina revealed robust expression of all four molecules in postnatal stages. In the postnatal retina, release of ACh by SACs is required for proper propagation of spontaneous activity in the form of retinal waves [40, 53]. Direction-selectivity in the retina, however, seems to emerge independently of visual experience and cholinergic waves and thus most likely arises due to complex molecular interactions [54]. Hence, the four ECM molecules are promising candidates that might contribute to the fine-tuning and establishment of neuronal circuits.

We observed that quadruple KO mice exhibit reduced a- and b-wave amplitudes, suggesting retinal dysfunction. We previously noted normal ERG responses in *Tnc* KO mice [55, 56]. Thus, we propose that the combined loss of four ECM molecules or alternatively the combined or single loss of Bcan, Ncan and Tnr may contribute to impaired ERG responses and retina function. Interestingly, we found that b-wave responses derived from excitatory glutamatergic rod BCs were most severely affected in the quadruple mutant. Cholinergic feedback to BCs enhances direction-selective signaling in postsynaptic SACs and DSGCs [57]. Therefore, the loss of cholinergic SACs and the reduced cholinergic feedback to rod BCs could explain the severe impairment of b-wave responses. The effects of the loss of ECM molecules on retinal functionality of sensory tissues such as the retina have not yet been described. So far, the ECM in tissues is rather known for its role in homeostasis and the organization of synapses.

We noted an imbalance of inhibitory GABAergic and excitatory glutamatergic signaling in the quadruple KO retina *in vivo*. Previous studies documented that the elimination of the four ECM proteins alters the ratio of excitatory and inhibitory hippocampal synapses *in vitro* [27]. Depletion experiments indicate that the ECM preserves the balanced state of neuronal networks by stabilizing inhibitory synapses [1]. Of note, an imbalance of GABA/glutamate has been suggested in albino mice with missing optokinetic nystagmus [58, 59]. Our findings suggest that mutual interactions of the matrisome regulate the delicate balance between GABAergic inhibition, glutamatergic excitation and visual motion processing *in vivo*. Tnc and Tnr bind Ncan and Bcan, respectively, with a high affinity [60, 61]. In astrocytes, Tnr modulates the uptake of glutamate [62]. A reduced GABAergic inhibition was found in the hippocampus after ablation of Tnr [19]. Based on these reports, we hypothesize that the cooperation of four matrix molecules is crucial for a balanced synaptic integrity. In that context, Tnr appears to exert a regulatory influence on GABAergic inhibition. Recently, it was reported that the neural ECM preserves the equilibrium of neuronal networks by stabilizing inhibitory synapses [29]. In this perspective, the reduction of the ECM might directly contribute to a weakening of inhibitory synapses.

Our MEA measurements revealed reduced DSGC responses, which reflect a defective visual processing in the quadruple KO retina. However, this did not reach statistical significance, which might be explained by the limited sensitivity of MEA measurements and/or compensatory mechanisms. In this regard, it is worth considering that various other ECM constituents, including laminins, collagens, several ECM receptors and ECM-modifying molecules were found dysregulated in our NGS analyses. Furthermore, the loss of SACs might be compensated by remaining SACs, which provide sufficient input to DSGCs.

In the CNS, tenascins exert a strong influence on synaptic structure and function [4, 5]. Thus, the absence of Tnc and Tnr in the quadruple KO could be one cause of impaired visual processing. To trace these limitations back to individual Tnc or Tnr genes, OMR measurements were carried out in single KO mice. When measured at both low and high velocities, *Tnc* KO mice showed a significantly reduced number of saccades compared to the WT. In contrast, in the *Tnr* KO the number of saccades was comparable to the WT at low velocities in either direction. At higher velocities, however, the number of saccades was also reduced in the *Tnr* KO. At slow velocities counterclockwise, a significant reduction of saccades was observed in *Tnc* KO in comparison to *Tnr* KO mice. These results indicate that the elimination of Tnc leads to limitations in visual processing at low and high velocities, whereas the *Tnr* deficiency mitigates processing only at high velocities, possibly due to compensatory effects. Yet, both the ablation of Tnc and Tnr impaired visual motion processing. The phenotype was accentuated in quadruple KO mice, where saccades were significantly diminished in contrast to both *Tnc* and *Tnr* KO mice. This strongly suggests that the loss of four ECM molecules intensified the limitations of visual motion processing that appeared to a milder degree in the single KOs. This may reflect gene-dosage effects of the matrisome and reveal cooperative synergism of the four ECM molecules. This may be based on molecular interactions as well as remote impact on gene regulation in the KO lines. The establishment of the precise pathways involved will require further investigations.

The ablation of SACs has been shown to decrease direction selectivity in retinal ON-OFF DSGCs *in vivo* [63]. ON-OFF DSGCs primarily project to the lateral geniculate nucleus (LGN) and the primary visual cortex (area V1 for visual motion integration [64–67]. Thus, it is reasonable to assume that deficiency of optomotor behavior observed in quadruple KO mice could potentially be explained by an impaired (sub-) cortical processing. In our recent study, we reported on PNN-associated ECM changes in the quadruple KO V1, which could directly affect optomotor behavior [28]. Notably, a distinct PNN-associated ECM expression was noted in the LGN, which may also play a role in visual motion processing [68]. Therefore, it is possible that cumulative changes in the ECM across various brain regions critical for visual motion computation could additionally contribute to the optomotor behavior deficiency observed in the quadruple KO model.

The causes for the restrictions of visual movement processing could also be disturbances in the synaptic circuitry and in the excitation/inhibition ratio of the retina. Interestingly, the NGS analyses are in accordance with the hypothesis of a synaptic imbalance in the quadruple KO retina, as a dysregulation of several synaptic candidates was registered. For example, we noted a downregulation of the RNA-binding protein *Rbpms2/Hermes*, a crucial modulator of synapse density as well as axon arbor formation in RGCs, which also influences optomotor processing [46]. We observed a downregulation of the *Kcnq2* gene, which is involved in the proper function of potassium channels in the brain. The functional importance of *Kcnq2* becomes particularly obvious under pathological conditions. Thus, alterations of KCNQ2 have been associated with seizures, autism as well as cognitive and developmental disabilities [69]. In a recent study, *Kcnq2* was presented as an important downstream regulator of ketamine in hippocampal glutamatergic neurons [70]. Also, we noted a downregulation of the G-protein coupled receptor *Glp1r* (*glucagon-like peptide-1 receptor*). The *Glp1r* agonist exendin-4 suppresses GABAR-mediated light-evoked inhibitory postsynaptic currents in RGCs [71]. We also found a downregulation of the *vesicular glutamate transporter 3/vGlut3 (Slc17a8).* Interestingly, vGlut3-expressing ACs provide excitatory input to ON-OFF and ON DSGCs and a subpopulation of W3 RGCs, but not SACs [72]. Therefore, the ECM loss in quadruple mutants might also impact SAC-independent excitatory glutamatergic signaling. In this regard, we observed downregulation of the *metabotropic glutamate receptor 1/Grm1* [73, 74]. Furthermore, we noted a downregulation of *Lrp1*, an important modulator of the ECM [75]. Interestingly, deletion of *Lrp1* from astrocytes in a hippocampal neuron co-culture model decreased neuronal network activity and influenced the proportion of pre- and postsynaptic structures [76]. In summary, we noted an extensive ECM remodeling and severe synaptic imbalance in quadruple mutant mice, which contribute to the impaired visual function and optomotor behavior.

A groundbreaking discovery had revealed that CSPG degradation by injection of chondroitinase-ABC into the visual cortex of adult rats leads to a reactivation of ocular dominance plasticity [77]. This observation provided evidence that the mature ECM inhibits experience-dependent plasticity and plays a central role in regulating visual input processing in the visual cortex. Conditions of degeneration such as in glaucoma are associated with complex ECM remodeling in the retina [30, 55, 56]. However, these studies mainly focused on structural ECM changes concurrent with visual impairment caused by disease, rather than on the direct effect of ECM on visual function. Hence, our study provides the first evidence for an impairment of sensorimotor integration in a neuronal subsystem as direct consequence of ECM deficiency.

### Limitations of the study

A limitation of our study is that the quadruple KO model provides insight into the cooperative effects of several ECM proteins but does not circumscribe the individual roles of the singular ECM molecule regarding the observed deficits in retina function. However, we revealed, yet less pronounced, optomotor deficits in single mutants for Tnc and Tnr. The relative contributions of the CSPGs Bcan and Ncan as well as the establishment of the pathways involved, will require further investigations.

## STAR METHODS

### Key resources table

| REAGENT or RESOURCE | SOURCE | IDENTIFIER |
| --- | --- | --- |
| Antibodies |  |  |
| Anti-Bcan | [78] | N/A |
| Anti-Brn3a | Santa Cruz Biotechnology | Cat# sc-31984;<br>RRID:AB_2167511 |
| Anti-ChAT | Millipore | Cat# AB144;<br>RRID:AB_11212843 |
| Anti-DIG Fab fragment antibody, AP-conjugated | Roche Diagnostics GmbH | Cat# 11093274910;<br>RRID:AB_514497 |
| Anti-GABA<sub>A</sub> receptor $\alpha 2$ | Synaptic Systems GmbH | Cat# 224 103;<br>RRID:AB_2108839 |
| Anti-Ncan | [79] | N/A |
| Anti-PSD95 | Millipore | Cat# MAB1598;<br>RRID:AB_94278 |
| Anti-Tnc | [80] | N/A |
| Anti-Tnr | [8] | N/A |
| Anti-VGLUT1 | Sigma-Aldrich | Cat# V0389;<br>RRID:AB_261840 |
| Alexa Fluor 488-AffiniPure Donkey Anti-Goat IgG (H+L) | Jackson ImmunoResearch Labs | Cat# 705-545-147;<br>RRID:AB_2336933 |
| Alexa Fluor 488-AffiniPure Goat Anti-Mouse IgG + IgM (H+L) | Jackson ImmunoResearch Labs | Cat# 115-545-044;<br>RRID:AB_2338844 |
| Alexa Fluor 488-AffiniPure Goat Anti-Rabbit IgG (H+L) | Jackson ImmunoResearch Labs | Cat# 111-545-045;<br>RRID:AB_2338049 |
| Biotin-SP-AffiniPure Goat Anti-Guinea Pig IgG (H+L) | Jackson ImmunoResearch Labs | Cat# 106-065-003;<br>RRID:AB_2337410 |
| Bacterial and virus strains |  |  |
| One Shot™ TOP10 chemically competent E. coli cells | Thermo Fisher Scientific | Cat# C404003 |
| Biological samples |  |  |
| WT and quadruple KO retina tissue | This work | N/A |
| Chemicals, peptides, and recombinant proteins |  |  |
| BamHI | Thermo Fisher Scientific | Cat# ER0051 |
| BCIP | Roche Diagnostics GmbH | Cat# 11383221001 |
| Blocking reagent | Roche Diagnostics GmbH | Cat# 11096176001 |
| Cy2-Streptavidin | Jackson ImmunoResearch Labs | Cat# 016-220-084 |
| EcoRI | Thermo Fisher Scientific | Cat# ER0271 |
| HindIII | Thermo Fisher Scientific | Cat#ER0501 |
| NBT | Roche Diagnostics GmbH | Cat# 11383213001 |
| TO-PRO™-3 iodide | Thermo Fisher Scientific | Cat# T3605 |
| XhoI | Thermo Fisher Scientific | Cat# ER0691 |
| Critical commercial assays |  |  |
| cDNA synthesis kit | Fermentas GmbH | Cat# K1612 |
| DIG RNA Labeling Mix | Roche Diagnostics GmbH | Cat# 26591020 |
| DirectPCR® Lysis Reagent Tail | VWR | Cat# 31-101-T |
| DNaseI (Rnase-free) | Roche Diagnostics GmbH | Cat# 04716728001 |
| JumpStart™ Tag DNA polymerase | Sigma-Aldrich | Cat# D9307 |
| NEBNext Ultra II Directional Total RNA Library Prep Kit for Illumina | NEB | Cat# E7760S/L |
| Proteinase K | Roth | Cat# 7528.3 |
| RNeasy® Mini Kit | Qiagen | Cat# 74104 |
| RNeasy® Midi Kit | Qiagen | Cat# 12143 |
| SP6 RNA polymerase | Roche Diagnostics GmbH | Cat# EP0131 |
| T7 RNA polymerase | Roche Diagnostics GmbH | Cat# EP0111 |
| TOPO® TA Cloning® Kit | Thermo Fisher Scientific | Cat# 45 - 0640 |
| QIAEX II® Gel Extraction Kit | Qiagen | Cat# 20021 |
| Experimental models: Cell lines |  |  |
| N/A | N/A | N/A |
| Experimental models: Organisms/strains |  |  |
| 129S2/SvPasCrl mice | Charles River Laboratories | RRID:IMSR_CRL:287 |
| Tnc KO mice | [81]<br>[82] | N/A |
| Tnr KO mice | [82] | N/A |
| Quadruple KO mice | [24] | N/A |
| Oligonucleotides |  |  |
| For primer pairs, see Tables S3 and S4 | Sigma-Aldrich | Tables S3 and S4 |
| Recombinant DNA |  |  |
| pCRTM2.1-TOPO vector | Thermo Fisher Scientific | Cat# 45 - 0640 |
| Software and algorithms |  |  |
| Adobe Illustrator | Adobe | http://www.adobe.com/products/illustrator.html; RRID:SCR_010279 |
| Adobe Photoshop | Adobe | https://www.adobe.com/products/photoshop.html; RRID:SCR_014199 |
| AxioVision Imaging System | Carl Zeiss | RRID:SCR_002677 |
| Black ZEN software | Carl Zeiss | RRID:SCR_018163 |
| Coral Paint Shop Pro | Coral Corporation | N/A |
| DESeq2 v.1.32.0 framework | [83] | https://bioconductor.org/packages/release/bioc/html/DESeq2.html; RRID:SCR_015687 |
| ERGView 4.380R | OcuScience | N/A |
| ImageJ | National Institutes of Health | https://imagej.nih.gov/ij/index.html; RRID:SCR_003070 |
| MATLAB | The MathWorks | http://www.mathworks.com/products/matlab/RRID:SCR_001622 |
| Offline Sorter | Plexon Inc | http://www.plexon.com/products/offline-sorter; RRID:SCR_000012 |
| R library gprofiler2 | [84] | https://biit.cs.ut.ee/gprofiler/page/r; RRID:SCR_018190 |
| Salmon v1.5.2 | [85] | https://combine-lab.github.io/salmon/; RRID:SCR_017036 |
| STATISTICA V13.3 | StatSoft | RRID:SCR_014213 |
| Other |  |  |
| 60MEA200/30IR-ITO | MultiChannel Systems | N/A |
| Axioplan2 | Carl Zeiss | N/A |
| BioSpectrometer | Eppendorf | N/A |
| CM3050 S cryostat | Leica | N/A |
| HMsERG system | OcuScience | N/A |
| Illumina NextSeq500 | Illumina | N/A |
| LSM 510 META | Carl Zeiss | N/A |
| TonoLab | Icare | N/A |
| Quantus Fluorometer | Promega | N/A |

### Experimental model and subject details

#### Animals

Mouse breeding colonies were maintained at the animal facility of the Faculty of Biology and Biotechnology, Ruhr University Bochum. Animals were housed under environmentally controlled lighting conditions (12-hour light-dark cycle) with access to chow and water *ad libitum*. Adult quadruple KO mice, generated by Rauch and colleagues through cross-breeding of the described single KO mutant mouse lines, combining the transgenic KO constructs of *Bcan, Ncan, Tnc* and *Tnr* [9, 17, 24, 81, 86] as well as mice only carrying the *Tnc* KO construct or the *Tnr* KO construct were compared to WT mice from the inbred background strain 129S2/SvPasCrl (Charles River Laboratories, Sulzfeld, Germany). Animals were mated overnight, and females were checked in the morning for the presence of a vaginal plug, which corresponded to embryonic day 0.5 (E0.5). For analyses, animals from the postnatal (P) stages P0, P4, P8 and P12 as well as adult mice of both sexes were euthanized by cervical decapitation or dislocation.

#### Animal study approval

The animal studies conducted in this work followed the guidelines of the Association for Research in Vision and Ophthalmology (ARVO) statement for the use of animals in ophthalmic and vision research. The experiments were performed in compliance with the German law (§15 TierSchG) and approved by the animal protection committee (Landesamt für Natur, Umwelt und Verbraucherschutz, Recklinghausen, North Rhine-Westphalia Germany; file number 84-02.04.2013.A291). The study was supervised by the animal welfare commissioner of the Ruhr University Bochum and the clinics facility for animal welfare (Einrichtung für Tierschutz, Tierärztlichen Dienst und Labortierkunde) of the University Tübingen. All efforts were made to minimize the number of animals used and their suffering.

### Method details

#### Genotyping

For the isolation of genomic DNA, mouse tail biopsies were lysed overnight at 55^°^C in DirectPCR^®^ Lysis Reagent Tail (VWR, Radnor, PA USA) containing 0.2 mg/ml proteinase K (Roth, Karlsruhe, Germany). The following day, samples were incubated for 45 minutes at 55^°^C. For genotyping, PCR analyses were performed using JumpStart^TM^ Taq DNA polymerase and primer pairs (Sigma-Aldrich, St. Louis, MO, USA) listed in Table S3. The following PCR conditions were used: 94°C for 2 minutes 40 seconds (initial denaturation), 37 cycles of 94°C for 30 seconds (denaturation), 56-60°C for 30 seconds (annealing) and 72°C for 50 seconds followed by 72°C for 5 minutes (final elongation). Following agarose gel electrophoresis (gel electrophoresis unit pharmacia biotech; Amersham, Freiburg, Germany) samples were documented under UV light in a gel documentation system (LFT Labortechnik GmbH & Co.KG, Wasserburg, Germany).

#### Intraocular pressure measurements

Intraocular pressure (IOP) measurements in quadruple KO and WT mice (16 weeks of age, both sexes) were performed with a rodent rebound tonometer (TonoLab; Icare; Vantaa, Finland) as previously described [87]. Both eyes were measured, and 10 readings of each eye were averaged. Data (N = 12) were compared by Student′s *t*-test (STATISTICA, V13.3; StatSoft, Hamburg, Germany) and are presented as mean ± standard error of the mean (SEM).

#### Electroretinogram recordings

Scotopic full-field flash electroretinogram (ERG) recordings were done with a HMsERG system (OcuScience, Henderson, NV, USA) as described previously [87]. Quadruple KO and WT mice of both genders (16 weeks of age) were dark-adapted overnight. For measurements, mice were hold under dim red light and anesthetized by an intraperitoneal injection of a ketamine-xylazine cocktail (120/16 mg/kg body weight). Afterward, eyes were topically anesthetized with Oxybuprocaine hydrochloride (Novesine Stulln^®^ 4 mg/ml, Stulln pharma, Stulln, Germany) and the pupil was dilated with Tropicamide (Mydriaticum Stulln^®^ 5 mg/ml, Pharma Stulln, Stulln, Germany). Scotopic flash series were recorded at 0.1, 0.3, 1.0, 3.0, 10 and 25 cd/m^2^. Then, electrical potentials were analyzed by the ERGView 4.380R software (OcuScience). A 150 Hz low-pass filter was applied, and a/b-wave amplitudes were evaluated. The a-wave amplitudes were measured from the pre-stimulus baseline up to the a-wave, while the b-waves were measured from the a-wave to the b-wave peak. Data (N = 8) were compared with STATISTICA (V13.3; StatSoft) using one-way analysis of variance (ANOVA) followed by Tukey′s post-hoc test and are shown as mean ± SEM.

#### Measurement of optomotor responses

To induce the optomotor response (OMR), mice were placed in a rotating cylinder (30 cm ø) with a vertical black and white stripe pattern. The stripes were 2 cm wide. The experiments were carried out in a defined order of stimulation: 1 minute cylinder rotation in one direction, 1 minute rest, 1 minute rotation in the other direction, 1 minute rest, repetition of the procedure at a changed velocity. Optimal velocities to elicit the OMR were 20^°^/second and 50^°^/second and rotation was done clockwise (CW) and counterclockwise (CCW). OMR was assessed by counting the fast-returning movement of the head after following the stripe pattern (defined as saccade). Analyses were done in male and female WT and quadruple KO as well as *Tnc* and *Tnr* single KO mice at 16 weeks of age. Data (N = 6) were compared with STATISTICA (V13.3; StatSoft) using one-way ANOVA followed by Tukey′s post-hoc test and are shown as mean ± SEM.

#### Multielectrode array

Microelectrode array (MEA) recordings have been reported previously [88]. To record spiking neural activity of RGCs, dark-adapted quadruple KO and WT retinae were prepared and mounted RGC side down contacting the 60-channel MEA (60MEA200/30IR-ITO; MultiChannel Systems, Reutlingen Germany). MEA-mounted retinae were superfused with artificial cerebrospinal fluid (ACSF) solution (125 mM NaCl, 2.5 mM KCl, 2 mM CaCl_2_, 1 mM MgCl_2_, 1.25 NaH_2_PO_4_, 26 mM NaHCO_3_ and 20 mM glucose) at a rate of 2.5 ml/minute at 33°C and left to recover for at least 30 minutes before recording. Extracellular voltage signals were recorded with equipment from MultiChannel Systems, processed offline to isolate spike trains using Offline Sorter (Plexon Inc., Dallas, TX, USA) and analyzed with custom scripts in MATLAB (The MathWorks, Natick, MA, USA). A visual stimulation set, including moving bars, was adapted from Baden and colleagues [89]. Stimuli were designed and delivered through QDSpy. The moving bar stimulus was represented as a white bar (300 x 1000 µm) drifting along its long axis in a sequence of positions covering the MEA and repeated at 8 different directions in sequence to quantify direction-selectivity. MEA data of quadruple KO and WT mice (N = 10) were analyzed in MATLAB (The MathWorks) and are shown as mean ± SEM.

#### Tissue collection for RT-PCR, immunohistochemistry and *in-situ* hybridization

For RNA extraction, retinae were isolated, pooled, snap frozen in liquid nitrogen and stored at −80°C until RNA extraction. For immunohistochemistry and *in-situ* hybridization of retinal sections, whole embryo heads, postnatal brains and adult eyes were fixed in 4% paraformaldehyde (PFA; Sigma-Aldrich) for 12-24 hours at 4°C and cryo-protected in 30% sucrose (J. T. Baker, Deventer, NL). Tissue was embedded in Jung TISSUE FREEZING MEDIUM^TM^ (Leica Instruments GmbH, St. Leon-Rot, Germany), sectioned in horizontal planes of 16 µm using a cryostat (CM3050 S, Leica, Bensheim, Germany) and collected onto Superfrost plus object slides (Menzel-Gläser, Braunschweig, Germany). Thereafter, slides were stored at −80°C until further processing. For the preparation of retinal flat-mounts, eyes were enucleated and fixed in 4% PFA at 4°C for 1 hour. Then, the retinae were dissected, post-fixed in 4% PFA for 5 minutes and washed in 1 x phosphate buffered saline (1 x PBS) until immunohistochemical staining.

#### RNA preparation, cDNA synthesis and reverse-transcription polymerase chain reaction

Total RNA was extracted using the RNeasy^®^ Mini or Midi kit (Qiagen, Hilden, Germany) and digested with Rnase-free DNaseI (Roche, Mannheim, Germany) according to the manufacturer’s instructions. The quality and quantity of RNA was assessed photometrically using the BioSpectrometer® (Eppendorf, Hamburg, Germany). To obtain cDNA, 1 µg of RNA was used for reverse transcription with a cDNA synthesis kit and random hexamer primers (Fermentas GmbH, St. Leon-Rot, Germany). Reverse-transcription polymerase chain reaction (RT-PCR) was performed using 1 µl cDNA using Taq DNA polymerase (JumpStart^TM^ Taq DNA polymerase (Sigma-Aldrich) in a Mastercycler Gradient (Eppendorf, Hamburg, Germany) with primer pairs listed in Table S4 (Sigma-Aldrich). The RT-PCR conditions were as follows: 94°C for 2 minutes 30 seconds (initial denaturation), 25-35 cycles of 94°C for 30 seconds (denaturation), 60°C for 45 seconds (annealing) and 72°C for 60 seconds followed by 72°C for 5 minutes (final elongation). Samples were documented as described above.

#### Next generation sequencing and bioinformatic analyses

For next generation sequencing (NGS), total RNA was isolated from adult WT and quadruple KO retinae (N = 4) as described above. The quantity of RNA was analyzed with the Quantus Fluorometer (Promega, Madison, WI, USA). RNA quality control was done with the RNA ScreenTape assay using the 4200 TapeStation system (Agilent Technologies Inc., Santa Clara, CA, USA). An RNA Integrity Number (RIN) of at least 7.8 verified the high quality of all included RNA samples. Sequencing libraries were generated from 450 ng of total RNA using the NEBNext Ultra II Directional Total RNA Library Prep Kit for Illumina as described by the manufacturer (NEB, Ipswich, MA, USA). The libraries were run on an Illumina NextSeq500 platform using the High Output 150 cycles Kit (Illumina, San Diego, CA, USA). FASTQ files were generated using bcl2fastq (Illumina). To facilitate reproducible analyses, samples were processed using the publicly available nf-core/RNA-seq pipeline version 3.5 [90] implemented in Nextflow 21.10.6 [91] using Docker 20.10.12 (Merkel 2014) with the minimal command. In brief, lane-level reads were trimmed with Trim Galore 0.6.7 [92] and aligned to the mouse genome (GRCm39) using STAR 2.7.9a [93]. Gene-level and transcript-level quantification was done by Salmon v1.5.2 [85]. Differential expression analyses were performed using custom scripts in R version 4.1.1 with the DESeq2 v.1.32.0 framework [83]. Genes were considered differentially expressed if the adjusted p-value was < 0.05. The 142 downregulated genes and 121 upregulated genes were further implemented in a gene ontology analyses with R library gprofiler2 [84] using gost function, correction_method = “fdr” and significant = TRUE. Gene ontology terms were considered significant with a false discovery rate < 0.05. As sources, we used Gene Ontology (GO or by branch GO:MF, GO:BP, GO:CC), Kyoto Encyclopedia of Genes and Genomes (KEGG), Reactome (REAC), WikiPathways (WP) and miRTarBase (MIRNA). The volcano plot and heatmaps were constructed with the R package ggplot2. The Integrative Genomics Viewer was used for genomic analyses of quadruple KO alleles [94] (Figure S5).

#### *In-situ* hybridization

The *in-situ* hybridization (ISH) procedure was performed following a protocol described by N. P. Pringle and W. D. Richardson (Wolfson Institute for Biomedical Research, London, UK). For the design of *in-situ* riboprobes, cDNAs of *Bcan*, *Ncan*, *Tnc* and *Tnr* were amplified with the ISH primers described above. Using the TOPO^TM^ TA Cloning^TM^ Kit with One Shot^TM^ TOP10 chemically competent E. coli cells (Thermo Fisher Scientific, Invitrogen, Waltham, MA, USA), gel purified (QIAEX II^®^ Gel Extraction Kit, Qiagen, Hilden Germany) RT-PCR products were cloned into the pCRTM2.1-TOPO vector (Thermo Fisher Scientific, Invitrogen) containing promotors for SP6 and T7 RNA polymerases (Roche Diagnostics GmbH, Mannheim, Germany). The generation of plasmids for *Tnc* and *Tnr* riboprobes was described previously [82]. After linearization with the respective restriction enzymes BamHI, EcoRI, HindIII or XhoI (Thermo Fisher Scientific, Waltham, MA, USA), digoxigenin (DIG)-labeled antisense and control sense RNA probes were made by *in vitro* transcription using the DIG RNA Labeling Mix (Roche Diagnostics GmbH) following the manufacturer’s protocol. For hybridization, DIG-labeled RNA probes were diluted 1:1,000 in Rnase-free hybridization buffer (1 x salts, 50% formamide, 0.1 mg/ml yeast tRNA, 10% dextran sulfate and 1 x Denhardt′s) heated to 75°C for 10 minutes, applied to retinal slices and incubated in a humified chamber (2 x saline sodium citrate (SSC) and 50% formamide) at 65°C overnight. The next day, slices were washed twice in washing buffer containing 1 x SSC, 50% formamide and 0.1% Tween-20 at 65°C for 30 minutes. Then, sections were washed twice in MABT buffer (100 mM maleic acid, 150 mM NaCl and 0.1% Tween-20, pH 7.5) at room temperature for 5 minutes. After stringent washing, retinal sections were transferred in a humified (H_2_O) chamber and incubated in blocking solution containing 2% blocking reagent (Roche Diagnostics GmbH, Mannheim, Germany) and 10% heat inactivated sheep serum in MABT buffer for 1 hour at room temperature. Afterward, slices were treated with alkaline phosphatase (AP)-conjugated anti-digoxigenin (DIG) Fab fragment antibody (sheep, 1:1,500; Roche Diagnostics GmbH) in blocking solution at 4°C overnight. Following antibody incubation, slides were washed three times in MABT buffer for 10 minutes and incubated twice in pre-developing buffer containing 100 mM Tris, pH 9.8, 100 mM NaCl and 50 mM MgCl_2_ for 15 minutes at room temperature. Hybridization signals were visualized by applying NBT/BCIP (4-Nitro blue tetrazolium chloride/1.5-bromo-4-chloro-3-indolyl-phophate; 1 mg/ml; Roche Diagnostics GmbH) developing solution (pre-developing solution and 5% polyvinyl alcohol, pH 9.5) for 2-4 hours at 37°C. To finish the staining reaction, retinal slides were washed in H_2_O, mounted with ImmuMount (Thermo Scientific Shandon^TM^, Waltham, MA, USA) and stored at 4°C.

#### Immunohistochemistry

Slides with retinal cross-sections were washed in 1 x PBS and incubated in blocking solution containing 1-3% normal goat or donkey serum (Dianova GmbH, Hamburg, Germany), 1% bovine serum albumin (BSA; AppliChem GmbH, Darmstadt, Germany) and 0.5% Triton^TM^-X-100 (Sigma-Aldrich, St. Louis, MO, USA) in 1 x PBS at room temperature for 1 hour. Then, primary antibodies were diluted in blocking solution and incubated at room temperature overnight. The following primary antibodies were used in this study: anti-Bcan (guinea pig, 1:250, [78]), anti-ChAT (goat, 1:200, Millipore, Billerica, MA, USA), anti-GABA_A_ receptor α2 (rabbit, 1:400, Synaptic Systems GmbH, Göttingen, Germany), anti-Tnc (batch KAF14/1, rabbit, 1:300, [80], anti-Ncan (rabbit, 1:300, [79]), anti-postsynaptic density protein 95 (anti-PSD95; mouse, 1:200, Millipore), anti-Tnr (anti-restrictin, clone 23-14, mouse, 1:100, Rathjen et al., 1991) and anti-VGLUT1 (rabbit, 1:400, Sigma-Aldrich). Following further washing in 1 x PBS, sections were incubated in blocking solution without Triton^TM^-X-100 containing appropriate secondary antibodies (donkey anti-goat, 1:300, Alexa Fluor 488, Jackson ImmunoResearch, West Grove, PA, USA; goat anti-mouse, 1:250, Alexa Fluor 488, Jackson ImmunoResearch; goat anti-guinea pig, 1:500, Biotin-SP, Jackson ImmunoResearch; goat anti-rabbit, 1:300, Alexa Fluor 488, Jackson ImmunoResearch. Biotin-Streptavidin immunostaining was detected with Streptavidin Cy2 (1:1,000, Jackson ImmunoResearch) was used. TO-PRO^TM^-3 iodide (1:400; Thermo Fisher Scientific, Invitrogen) was added to visualize cell nuclei. Retinal flat-mounts were blocked in blocking solution containing 3% donkey serum, 1% BSA and 2% Triton^TM^-X-100 in 1 x PBS at room temperature for 1 hour. Thereafter, flat-mounts were incubated in blocking solution with antibodies directed against ChAT (Millipore) and Brn3a (brain-specific homeobox/POU domain protein 3a; goat, 1:300, Santa Cruz Biotechnology, Dallas, TX, USA) at 4°C for 48 hours. Flat-mounts were washed in 1 x PBS and incubated with donkey anti-goat secondary antibodies (1:300, Alexa Fluor 488, Jackson ImmunoResearch) and TO-PRO^TM^-3 iodide (1:400, Thermo Fisher Scientific, Invitrogen) in blocking solution without Triton^TM^-X-100 for 2 hours at room temperature.

For microscopic analyses, retinal cross-sections and flat-mounts were cover-slipped with ImmuMount (Thermo Scientific Shandon^TM^). For retinal section analyses, images were taken from 2 retinal sections per slide. 2 peripheral and 2 central images per retina at a 400 x magnification were captured. To examine VGLUT1 staining, images (IPL: 100 x 240 pixel, OPL: 100 x 70 pixel), were cropped with Coral Paint Shop Pro X8 (Coral Corporation, Ottawa, CA). ImageJ software (National Institutes of Health; Bethesda, MD, USA) was used to perform masked evaluation of the staining signal as described previously [87]. Images were converted into grey scales. The background was subtracted and lower (IPL: 17.7 pixel, OPL: 28.8 pixel) and upper threshold (IPL: 90.7 pixel, OPL: 86.4 pixel) values were set. Then, the percentage of the area fraction was measured as described [87]. Cell counting was done for immunostainings of ChAT and GABA_A_ receptor α2. To analyze cell numbers in retinal flat-mounts, immunoreactive cells were counted in the peripheral and central retina (200 x magnification; 8 counting windows/retinal flat-mount per animal; 2 counting windows from each retinal quadrant). To quantify immunohistochemical staining, data (N = 3-8) were compared with Student′s *t*-test (STATISTICA; V13.3; StatSoft).

#### Microscopy, image processing and statistical analyses

For the documentation of the *in-situ* hybridization, images were acquired using the Axioplan2 microscope equipped with an AxioCam MRm digital camera and the AxioVision 4.5 software (Carl Zeiss, Oberkochen, Germany). Fluorescence specimens were examined by confocal laser-scanning microscopy (LSM 510 META; Carl Zeiss). Laser lines and emission filters were adjusted using the Zeiss Black ZEN software (Carl Zeiss). Images were exported and processed with Adobe Photoshop and Adobe Illustrator (Adobe, Dublin, Ireland), Coral Paint Shop Pro X8 (Coral Corporation) and ImageJ (National Institutes of Health). Data are presented as mean ± SEM and/or ± standard deviation (SD) and groups were analyzed with Matlab (MathWorks) or Statistica (StatSoft) using Student′s *t*-test or one-way ANOVA followed by Tukey′s post-hoc test. Values of p < 0.05 were considered statistically significant.

## Supporting information

Supplement_Reinhard et al. 2023

## ACKNOWLEDGEMENTS

We are grateful to Constanze Seidenbecher, Uwe Rauch and Fritz Rathjen for the generous gift of anti-Bcan, anti-Ncan and anti-Tnr antibodies, respectively. We acknowledge Claudia Distler-Hoffmann and Hermann Korbmacher for their helpful advice concerning the optomotor response recordings. We thank Marina Palmhof for technical assistance on the electroretinogram measurements. We are grateful to Stephanie Chun, Klaudija Masarini, Anna Rudzinski and Marion Voelzkow for excellent technical support. This work was supported by the Genomics Facility, a core facility of the Interdisciplinary Center for Clinical Research (IZKF) Aachen within the Faculty of Medicine at RWTH Aachen University.

## AUTHOR CONTRIBUTIONS

Conceptualization, J.R. and A.F.; methodology, J.R., C.M., S.W., L.R., V.L., H.S., D.L.R., L.G., C.C.K., J.F., S.C.J and A.F.; formal analysis, J.R., C.M., S.W., L.R., V.L., H.S., D.L.R., L.G., C.C.K.; investigation, J.R., C.M., S.W., L.R., V.L., H.S., L.G. and C.C.K.; writing – original draft, J.R.; writing & editing, C.M., S.W., L.R., H.S., D.L.R., L.G., C.C.K., J.F., S.C.J. and A.F.; visualization, J.R., C.M., S.W., L.R., V.L., H.S., D.L.R., L.G. and C.C.K.; funding acquisition, S.W., V.L, A.F.

## REFERENCES

1. Ferrer-Ferrer M, Dityatev A: Shaping Synapses by the Neural Extracellular Matrix. Front Neuroanat 2018, 12:40.

2. Heikkinen A, Pihlajaniemi T, Faissner A, Yuzaki M: Neural ECM and synaptogenesis. Progress in brain research 2014, 214:29–51.

3. Dzyubenko E, Gottschling C, Faissner A: Neuron-Glia Interactions in Neural Plasticity: Contributions of Neural Extracellular Matrix and Perineuronal Nets. Neural Plast 2016, 2016:5214961.

4. Dityatev A, Schachner M, Sonderegger P: The dual role of the extracellular matrix in synaptic plasticity and homeostasis. Nat Rev Neurosci 2010, 11(11):735–746.

5. Fawcett JW, Oohashi T, Pizzorusso T: The roles of perineuronal nets and the perinodal extracellular matrix in neuronal function. Nat Rev Neurosci 2019, 20(8):451–465.

6. Evers MR, Salmen B, Bukalo O, Rollenhagen A, Bosl MR, Morellini F, Bartsch U, Dityatev A, Schachner M: Impairment of L-type Ca2+ channel-dependent forms of hippocampal synaptic plasticity in mice deficient in the extracellular matrix glycoprotein tenascin-C. J Neurosci 2002, 22(16):7177–7194.

7. Faissner A: The tenascin gene family in axon growth and guidance. Cell Tissue Res 1997, 290(2):331–341.

8. Rathjen FG, Hodge R: Early Days of Tenascin-R Research: Two Approaches Discovered and Shed Light on Tenascin-R. Front Immunol 2020, 11:612482.

9. Weber P, Bartsch U, Rasband MN, Czaniera R, Lang Y, Bluethmann H, Margolis RU, Levinson SR, Shrager P, Montag D et al: Mice deficient for tenascin-R display alterations of the extracellular matrix and decreased axonal conduction velocities in the CNS. J Neurosci 1999, 19(11):4245–4262.

10. Iozzo RV, Schaefer L: Proteoglycan form and function: A comprehensive nomenclature of proteoglycans. Matrix Biol 2015, 42:11–55.

11. Maeda N: Proteoglycans and neuronal migration in the cerebral cortex during development and disease. Frontiers in neuroscience 2015, 9:98.

12. Inatani M, Honjo M, Otori Y, Oohira A, Kido N, Tano Y, Honda Y, Tanihara H: Inhibitory effects of neurocan and phosphacan on neurite outgrowth from retinal ganglion cells in culture. Invest Ophthalmol Vis Sci 2001, 42(8):1930–1938.

13. Yamada H, Fredette B, Shitara K, Hagihara K, Miura R, Ranscht B, Stallcup WB, Yamaguchi Y: The brain chondroitin sulfate proteoglycan brevican associates with astrocytes ensheathing cerebellar glomeruli and inhibits neurite outgrowth from granule neurons. J Neurosci 1997, 17(20):7784–7795.

14. Milev P, Maurel P, Chiba A, Mevissen M, Popp S, Yamaguchi Y, Margolis RK, Margolis RU: Differential regulation of expression of hyaluronan-binding proteoglycans in developing brain: aggrecan, versican, neurocan, and brevican. Biochem Biophys Res Commun 1998, 247(2):207–212.

15. Deepa SS, Carulli D, Galtrey C, Rhodes K, Fukuda J, Mikami T, Sugahara K, Fawcett JW: Composition of perineuronal net extracellular matrix in rat brain: a different disaccharide composition for the net-associated proteoglycans. J Biol Chem 2006, 281(26):17789–17800.

16. Hagihara K, Miura R, Kosaki R, Berglund E, Ranscht B, Yamaguchi Y: Immunohistochemical evidence for the brevican-tenascin-R interaction: colocalization in perineuronal nets suggests a physiological role for the interaction in the adult rat brain. J Comp Neurol 1999, 410(2):256–264.

17. Brakebusch C, Seidenbecher CI, Asztely F, Rauch U, Matthies H, Meyer H, Krug M, Bockers TM, Zhou X, Kreutz MR et al: Brevican-deficient mice display impaired hippocampal CA1 long-term potentiation but show no obvious deficits in learning and memory. Mol Cell Biol 2002, 22(21):7417–7427.

18. Saghatelyan AK, Dityatev A, Schmidt S, Schuster T, Bartsch U, Schachner M: Reduced perisomatic inhibition, increased excitatory transmission, and impaired long-term potentiation in mice deficient for the extracellular matrix glycoprotein tenascin-R. Mol Cell Neurosci 2001, 17(1):226–240.

19. Bukalo O, Schachner M, Dityatev A: Modification of extracellular matrix by enzymatic removal of chondroitin sulfate and by lack of tenascin-R differentially affects several forms of synaptic plasticity in the hippocampus. Neuroscience 2001, 104(2):359–369.

20. Masland RH: The tasks of amacrine cells. Vis Neurosci 2012, 29(1):3–9.

21. Balasubramanian R, Gan L: Development of Retinal Amacrine Cells and Their Dendritic Stratification. Curr Ophthalmol Rep 2014, 2(3):100–106.

22. Bleckert A, Zhang C, Turner MH, Koren D, Berson DM, Park SJH, Demb JB, Rieke F, Wei W, Wong RO: GABA release selectively regulates synapse development at distinct inputs on direction-selective retinal ganglion cells. Proc Natl Acad Sci U S A 2018, 115(51):E12083–E12090.

23. Auferkorte ON, Baden T, Kaushalya SK, Zabouri N, Rudolph U, Haverkamp S, Euler T: GABA(A) receptors containing the alpha2 subunit are critical for direction-selective inhibition in the retina. PLoS One 2012, 7(4):e35109.

24. Rauch U, Zhou XH, Roos G: Extracellular matrix alterations in brains lacking four of its components. Biochem Biophys Res Commun 2005, 328(2):608–617.

25. Geissler M, Gottschling C, Aguado A, Rauch U, Wetzel CH, Hatt H, Faissner A: Primary hippocampal neurons, which lack four crucial extracellular matrix molecules, display abnormalities of synaptic structure and function and severe deficits in perineuronal net formation. J Neurosci 2013, 33(18):7742–7755.

26. Jansen S, Gottschling C, Faissner A, Manahan-Vaughan D: Intrinsic cellular and molecular properties of in vivo hippocampal synaptic plasticity are altered in the absence of key synaptic matrix molecules. Hippocampus 2017, 27(8):920–933.

27. Gottschling C, Wegrzyn D, Denecke B, Faissner A: Elimination of the four extracellular matrix molecules tenascin-C, tenascin-R, brevican and neurocan alters the ratio of excitatory and inhibitory synapses. Scientific reports 2019, 9(1):13939.

28. Mueller-Buehl C, Reinhard J, Roll L, Bader V, Winklhofer KF, Faissner A: Brevican, Neurocan, Tenascin-C, and Tenascin-R Act as Important Regulators of the Interplay Between Perineuronal Nets, Synaptic Integrity, Inhibitory Interneurons, and Otx2. Front Cell Dev Biol 2022, 10:886527.

29. Dzyubenko E, Fleischer M, Manrique-Castano D, Borbor M, Kleinschnitz C, Faissner A, Hermann DM: Inhibitory control in neuronal networks relies on the extracellular matrix integrity. Cell Mol Life Sci 2021, 78(14):5647–5663.

30. Al-Ubaidi MR, Naash MI, Conley SM: A perspective on the role of the extracellular matrix in progressive retinal degenerative disorders. Invest Ophthalmol Vis Sci 2013, 54(13):8119–8124.

31. Huxlin KR, Carr R, Schulz M, Sefton AJ, Bennett MR: Trophic effect of collicular proteoglycan on neonatal rat retinal ganglion cells in situ. Brain Res Dev Brain Res 1995, 84(1):77–88.

32. Inatani M, Tanihara H, Oohira A, Honjo M, Honda Y: Identification of a nervous tissue-specific chondroitin sulfate proteoglycan, neurocan, in developing rat retina. Invest Ophthalmol Vis Sci 1999, 40(10):2350–2359.

33. D’Alessandri L, Ranscht B, Winterhalter KH, Vaughan L: Contactin/F11 and tenascin-C co-expression in the chick retina correlates with formation of the synaptic plexiform layers. Curr Eye Res 1995, 14(10):911–926.

34. Bartsch U, Pesheva P, Raff M, Schachner M: Expression of janusin (J1-160/180) in the retina and optic nerve of the developing and adult mouse. Glia 1993, 9(1):57–69.

35. Kretschmer F, Tariq M, Chatila W, Wu B, Badea TC: Comparison of optomotor and optokinetic reflexes in mice. J Neurophysiol 2017, 118(1):300–316.

36. Yoshida K, Watanabe D, Ishikane H, Tachibana M, Pastan I, Nakanishi S: A key role of starburst amacrine cells in originating retinal directional selectivity and optokinetic eye movement. Neuron 2001, 30(3):771–780.

37. Amthor FR, Keyser KT, Dmitrieva NA: Effects of the destruction of starburst-cholinergic amacrine cells by the toxin AF64A on rabbit retinal directional selectivity. Vis Neurosci 2002, 19(4):495–509.

38. Rasmussen R, Matsumoto A, Dahlstrup Sietam M, Yonehara K: A segregated cortical stream for retinal direction selectivity. Nat Commun 2020, 11(1):831.

39. Famiglietti EV, Jr.: On and off pathways through amacrine cells in mammalian retina: the synaptic connections of “starburst” amacrine cells. Vision Res 1983, 23(11):1265–1279.

40. Bansal A, Singer JH, Hwang BJ, Xu W, Beaudet A, Feller MB: Mice lacking specific nicotinic acetylcholine receptor subunits exhibit dramatically altered spontaneous activity patterns and reveal a limited role for retinal waves in forming ON and OFF circuits in the inner retina. J Neurosci 2000, 20(20):7672–7681.

41. Zhang J, Yang Z, Wu SM: Development of cholinergic amacrine cells is visual activity-dependent in the postnatal mouse retina. J Comp Neurol 2005, 484(3):331–343.

42. Johnson J, Fremeau RT, Jr., Duncan JL, Renteria RC, Yang H, Hua Z, Liu X, LaVail MM, Edwards RH, Copenhagen DR: Vesicular glutamate transporter 1 is required for photoreceptor synaptic signaling but not for intrinsic visual functions. J Neurosci 2007, 27(27):7245–7255.

43. Johnson J, Tian N, Caywood MS, Reimer RJ, Edwards RH, Copenhagen DR: Vesicular neurotransmitter transporter expression in developing postnatal rodent retina: GABA and glycine precede glutamate. J Neurosci 2003, 23(2):518–529.

44. Sherry DM, Wang MM, Bates J, Frishman LJ: Expression of vesicular glutamate transporter 1 in the mouse retina reveals temporal ordering in development of rod vs. cone and ON vs. OFF circuits. J Comp Neurol 2003, 465(4):480–498.

45. Kim IB, Oh SJ, Nielsen S, Chun MH: Immunocytochemical localization of aquaporin 1 in the rat retina. Neurosci Lett 1998, 244(1):52–54.

46. Hornberg H, Wollerton-van Horck F, Maurus D, Zwart M, Svoboda H, Harris WA, Holt CE: RNA-binding protein Hermes/RBPMS inversely affects synapse density and axon arbor formation in retinal ganglion cells in vivo. J Neurosci 2013, 33(25):10384–10395.

47. Barlow HB, Hill RM, Levick WR: Retinal Ganglion Cells Responding Selectively to Direction and Speed of Image Motion in the Rabbit. J Physiol 1964, 173:377–407.

48. Taylor WR, Smith RG: The role of starburst amacrine cells in visual signal processing. Vis Neurosci 2012, 29(1):73–81.

49. Jones FS, Jones PL: The tenascin family of ECM glycoproteins: structure, function, and regulation during embryonic development and tissue remodeling. Dev Dyn 2000, 218(2):235–259.

50. Reinhard J, Joachim SC, Faissner A: Extracellular matrix remodeling during retinal development. Exp Eye Res 2015, 133:132–140.

51. Theocharis AD, Skandalis SS, Gialeli C, Karamanos NK: Extracellular matrix structure. Adv Drug Deliv Rev 2016, 97:4–27.

52. Reinhard J, Roll L, Faissner A: Tenascins in Retinal and Optic Nerve Neurodegeneration. Front Integr Neurosci 2017, 11:30.

53. Feller MB, Wellis DP, Stellwagen D, Werblin FS, Shatz CJ: Requirement for cholinergic synaptic transmission in the propagation of spontaneous retinal waves. Science 1996, 272(5265):1182–1187.

54. Elstrott J, Anishchenko A, Greschner M, Sher A, Litke AM, Chichilnisky EJ, Feller MB: Direction selectivity in the retina is established independent of visual experience and cholinergic retinal waves. Neuron 2008, 58(4):499–506.

55. Wiemann S, Yousf A, Joachim SC, Peters C, Mueller-Buehl AM, Wagner N, Reinhard J: Knock-Out of Tenascin-C Ameliorates Ischemia-Induced Rod-Photoreceptor Degeneration and Retinal Dysfunction. Frontiers in neuroscience 2021, 15:642176.

56. Wiemann S, Reinhard J, Reinehr S, Cibir Z, Joachim SC, Faissner A: Loss of the Extracellular Matrix Molecule Tenascin-C Leads to Absence of Reactive Gliosis and Promotes Anti-inflammatory Cytokine Expression in an Autoimmune Glaucoma Mouse Model. Front Immunol 2020, 11:566279.

57. Hellmer CB, Hall LM, Bohl JM, Sharpe ZJ, Smith RG, Ichinose T: Cholinergic feedback to bipolar cells contributes to motion detection in the mouse retina. Cell Rep 2021, 37(11):110106.

58. Blaszczyk WM, Telkes I, Distler C: GABA-immunoreactive starburst amacrine cells in pigmented and albino rats. Eur J Neurosci 2004, 20(11):3195–3198.

59. Blaszczyk WM, Straub H, Distler C: GABA content in the retina of pigmented and albino rats. Neuroreport 2004, 15(7):1141–1144.

60. Day JM, Olin AI, Murdoch AD, Canfield A, Sasaki T, Timpl R, Hardingham TE, Aspberg A: Alternative splicing in the aggrecan G3 domain influences binding interactions with tenascin-C and other extracellular matrix proteins. J Biol Chem 2004, 279(13):12511–12518.

61. Midwood KS, Orend G: The role of tenascin-C in tissue injury and tumorigenesis. Journal of cell communication and signaling 2009, 3(3-4):287–310.

62. Okuda H, Tatsumi K, Morita S, Shibukawa Y, Korekane H, Horii-Hayashi N, Wada Y, Taniguchi N, Wanaka A: Chondroitin sulfate proteoglycan tenascin-R regulates glutamate uptake by adult brain astrocytes. J Biol Chem 2014, 289(5):2620–2631.

63. Hillier D, Fiscella M, Drinnenberg A, Trenholm S, Rompani SB, Raics Z, Katona G, Juettner J, Hierlemann A, Rozsa B et al: Causal evidence for retina-dependent and -independent visual motion computations in mouse cortex. Nat Neurosci 2017, 20(7):960–968.

64. Priebe NJ, Ferster D: Direction selectivity of excitation and inhibition in simple cells of the cat primary visual cortex. Neuron 2005, 45(1):133–145.

65. Huberman AD, Wei W, Elstrott J, Stafford BK, Feller MB, Barres BA: Genetic identification of an On-Off direction-selective retinal ganglion cell subtype reveals a layer-specific subcortical map of posterior motion. Neuron 2009, 62(3):327–334.

66. Kay JN, De la Huerta I, Kim IJ, Zhang Y, Yamagata M, Chu MW, Meister M, Sanes JR: Retinal ganglion cells with distinct directional preferences differ in molecular identity, structure, and central projections. J Neurosci 2011, 31(21):7753–7762.

67. Rivlin-Etzion M, Zhou K, Wei W, Elstrott J, Nguyen PL, Barres BA, Huberman AD, Feller MB: Transgenic mice reveal unexpected diversity of on-off direction-selective retinal ganglion cell subtypes and brain structures involved in motion processing. J Neurosci 2011, 31(24):8760–8769.

68. Sabbagh U, Monavarfeshani A, Su K, Zabet-Moghadam M, Cole J, Carnival E, Su J, Mirzaei M, Gupta V, Salekdeh GH et al: Distribution and development of molecularly distinct perineuronal nets in visual thalamus. J Neurochem 2018, 147(5):626–646.

69. Miceli F, Soldovieri MV, Weckhuysen S, Cooper E, Taglialatela M: KCNQ2-Related Disorders. In: GeneReviews((R)). Edited by Adam MP, Everman DB, Mirzaa GM, Pagon RA, Wallace SE, Bean LJH, Gripp KW, Amemiya A. Seattle (WA); 1993.

70. Lopez JP, Lucken MD, Brivio E, Karamihalev S, Kos A, De Donno C, Benjamin A, Yang H, Dick ALW, Stoffel R et al: Ketamine exerts its sustained antidepressant effects via cell-type-specific regulation of Kcnq2. Neuron 2022.

71. Zhang T, Ruan HZ, Wang YC, Shao YQ, Zhou W, Weng SJ, Zhong YM: Signaling Mechanism for Modulation by GLP-1 and Exendin-4 of GABA Receptors on Rat Retinal Ganglion Cells. Neurosci Bull 2022.

72. Lee S, Chen L, Chen M, Ye M, Seal RP, Zhou ZJ: An unconventional glutamatergic circuit in the retina formed by vGluT3 amacrine cells. Neuron 2014, 84(4):708–715.

73. Haverkamp S, Wassle H: Characterization of an amacrine cell type of the mammalian retina immunoreactive for vesicular glutamate transporter 3. J Comp Neurol 2004, 468(2):251–263.

74. Li Q, Cui P, Miao Y, Gao F, Li XY, Qian WJ, Jiang SX, Wu N, Sun XH, Wang Z: Activation of group I metabotropic glutamate receptors regulates the excitability of rat retinal ganglion cells by suppressing Kir and I h. Brain Struct Funct 2017, 222(2):813–830.

75. Bres EE, Faissner A: Low Density Receptor-Related Protein 1 Interactions With the Extracellular Matrix: More Than Meets the Eye. Front Cell Dev Biol 2019, 7:31.

76. Romeo R, Glotzbach K, Scheller A, Faissner A: Deletion of LRP1 From Astrocytes Modifies Neuronal Network Activity in an in vitro Model of the Tripartite Synapse. Front Cell Neurosci 2020, 14:567253.

77. Pizzorusso T, Medini P, Berardi N, Chierzi S, Fawcett JW, Maffei L: Reactivation of ocular dominance plasticity in the adult visual cortex. Science 2002, 298(5596):1248–1251.

78. Seidenbecher CI, Richter K, Rauch U, Fassler R, Garner CC, Gundelfinger ED: Brevican, a chondroitin sulfate proteoglycan of rat brain, occurs as secreted and cell surface glycosylphosphatidylinositol-anchored isoforms. J Biol Chem 1995, 270(45):27206–27212.

79. Haas CA, Rauch U, Thon N, Merten T, Deller T: Entorhinal cortex lesion in adult rats induces the expression of the neuronal chondroitin sulfate proteoglycan neurocan in reactive astrocytes. J Neurosci 1999, 19(22):9953–9963.

80. Faissner A, Kruse J: J1/tenascin is a repulsive substrate for central nervous system neurons. Neuron 1990, 5(5):627–637.

81. Forsberg E, Hirsch E, Frohlich L, Meyer M, Ekblom P, Aszodi A, Werner S, Fassler R: Skin wounds and severed nerves heal normally in mice lacking tenascin-C. Proc Natl Acad Sci U S A 1996, 93(13):6594–6599.

82. Czopka T, Von Holst A, Schmidt G, Ffrench-Constant C, Faissner A: Tenascin C and tenascin R similarly prevent the formation of myelin membranes in a RhoA-dependent manner, but antagonistically regulate the expression of myelin basic protein via a separate pathway. Glia 2009, 57(16):1790–1801.

83. Love MI, Huber W, Anders S: Moderated estimation of fold change and dispersion for RNA-seq data with DESeq2. Genome Biol 2014, 15(12):550.

84. Kolberg L, Raudvere U, Kuzmin I, Vilo J, Peterson H: gprofiler2 -- an R package for gene list functional enrichment analysis and namespace conversion toolset g:Profiler. F1000Res 2020, 9.

85. Patro R, Duggal G, Love MI, Irizarry RA, Kingsford C: Salmon provides fast and bias-aware quantification of transcript expression. Nat Methods 2017, 14(4):417–419.

86. Zhou XH, Brakebusch C, Matthies H, Oohashi T, Hirsch E, Moser M, Krug M, Seidenbecher CI, Boeckers TM, Rauch U et al: Neurocan is dispensable for brain development. Mol Cell Biol 2001, 21(17):5970–5978.

87. Reinhard J, Wiemann S, Joachim SC, Palmhof M, Woestmann J, Denecke B, Wang Y, Downey GP, Faissner A: Heterozygous Meg2 Ablation Causes Intraocular Pressure Elevation and Progressive Glaucomatous Neurodegeneration. Molecular neurobiology 2019, 56(6):4322–4345.

88. Jalligampala A, Sekhar S, Zrenner E, Rathbun DL: Optimal voltage stimulation parameters for network-mediated responses in wild type and rd10 mouse retinal ganglion cells. J Neural Eng 2017, 14(2):026004.

89. Baden T, Berens P, Franke K, Roman Roson M, Bethge M, Euler T: The functional diversity of retinal ganglion cells in the mouse. Nature 2016, 529(7586):345–350.

90. Ewels PA, Peltzer A, Fillinger S, Patel H, Alneberg J, Wilm A, Garcia MU, Di Tommaso P, Nahnsen S: The nf-core framework for community-curated bioinformatics pipelines. Nat Biotechnol 2020, 38(3):276–278.

91. Di Tommaso P, Chatzou M, Floden EW, Barja PP, Palumbo E, Notredame C: Nextflow enables reproducible computational workflows. Nat Biotechnol 2017, 35(4):316–319.

92. Krueger F, James F, Ewels P, Afyounian E, Schuster-Boeckler B: FelixKrueger/TrimGalore: V0.6.7 - DOI via Zenodo (version 0.6.7). Zenodo https://doiorg/105281/zenodo5127899 2021.

93. Dobin A, Davis CA, Schlesinger F, Drenkow J, Zaleski C, Jha S, Batut P, Chaisson M, Gingeras TR: STAR: ultrafast universal RNA-seq aligner. Bioinformatics 2013, 29(1):15–21.

94. Robinson JT, Thorvaldsdottir H, Winckler W, Guttman M, Lander ES, Getz G, Mesirov JP: Integrative genomics viewer. Nat Biotechnol 2011, 29(1):24–26.

95. Talts JF, Wirl G, Dictor M, Muller WJ, Fassler R: Tenascin-C modulates tumor stroma and monocyte/macrophage recruitment but not tumor growth or metastasis in a mouse strain with spontaneous mammary cancer. J Cell Sci 1999, 112 (Pt 12):1855–1864.

