## Supplement_Reinhard et al. 2023 for "Neural extracellular matrix regulates visual sensory motor integration"

**Suppl. figures and tables**

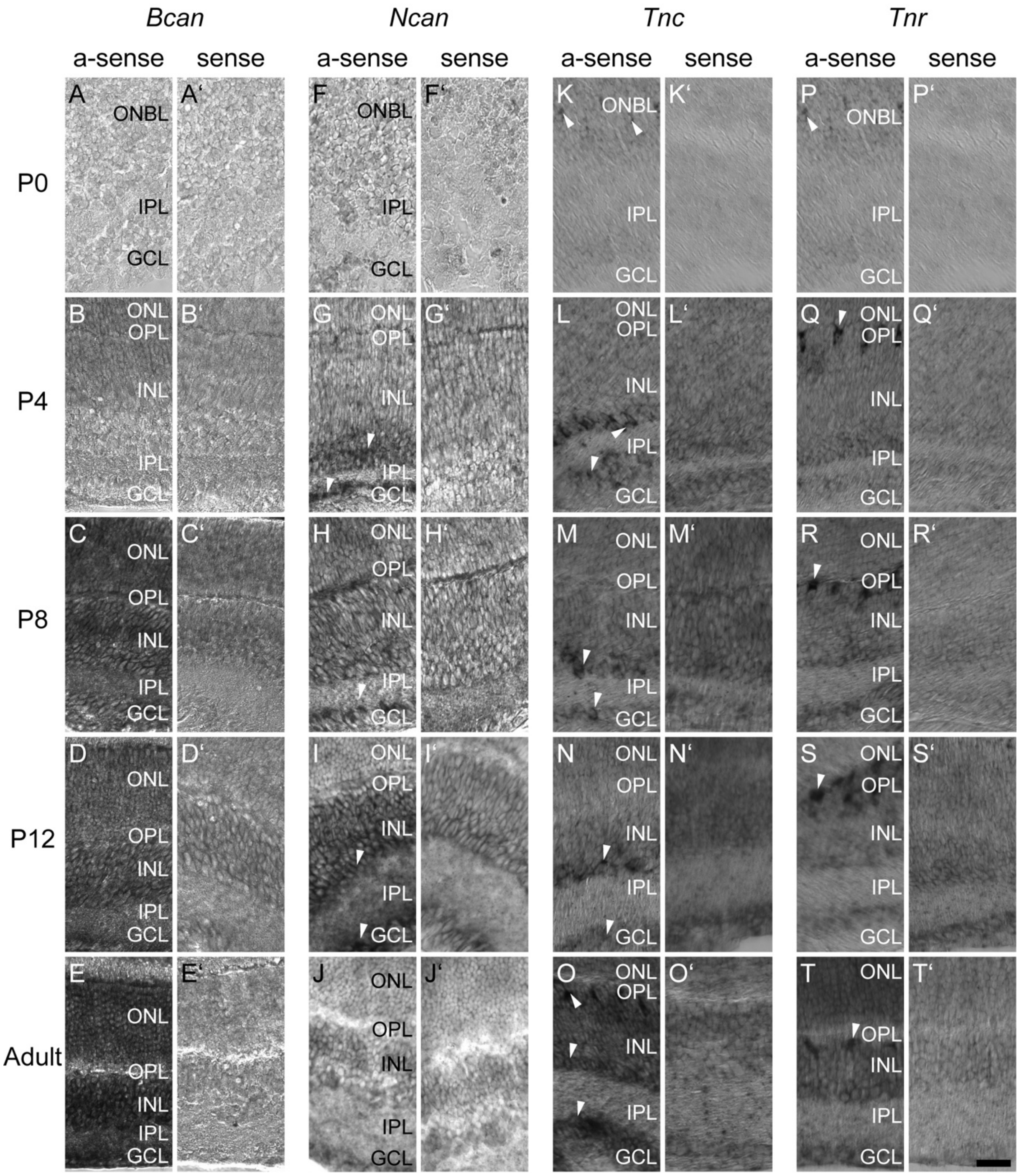


**Figure S1. *In-situ* hybridization revealing the spatiotemporal expression pattern of *Bcan*, *Ncan*, *Tnc* and *Tnr* in the postnatal and adult mouse retina.** Differential mRNA expression pattern of *Bcan* (A-E, anti-sense; A`-E`, sense), *Ncan* (F-J, anti-sense; F`-J`, sense), *Tnc* (K-O, anti-sense; K`-O`, sense) and *Tnr* (P-T, anti-sense; P`-T`, sense) in the postnatal (P0, P4, P8 and P12) and adult retina. White arrows mark prominent signals. Scale bar: 20 µm. *Bcan* = *brevican*; GCL = ganglion cell layer; INL = inner nuclear layer; IPL = inner plexiform layer; *Ncan* = *neurocan*; ONBL = outer neuroblastic layer; ONL = outer nuclear layer; OPL = outer plexiform layer; P = postnatal; *Tnc* = *tenascin-C*; *Tnr* = *tenascin-R*.

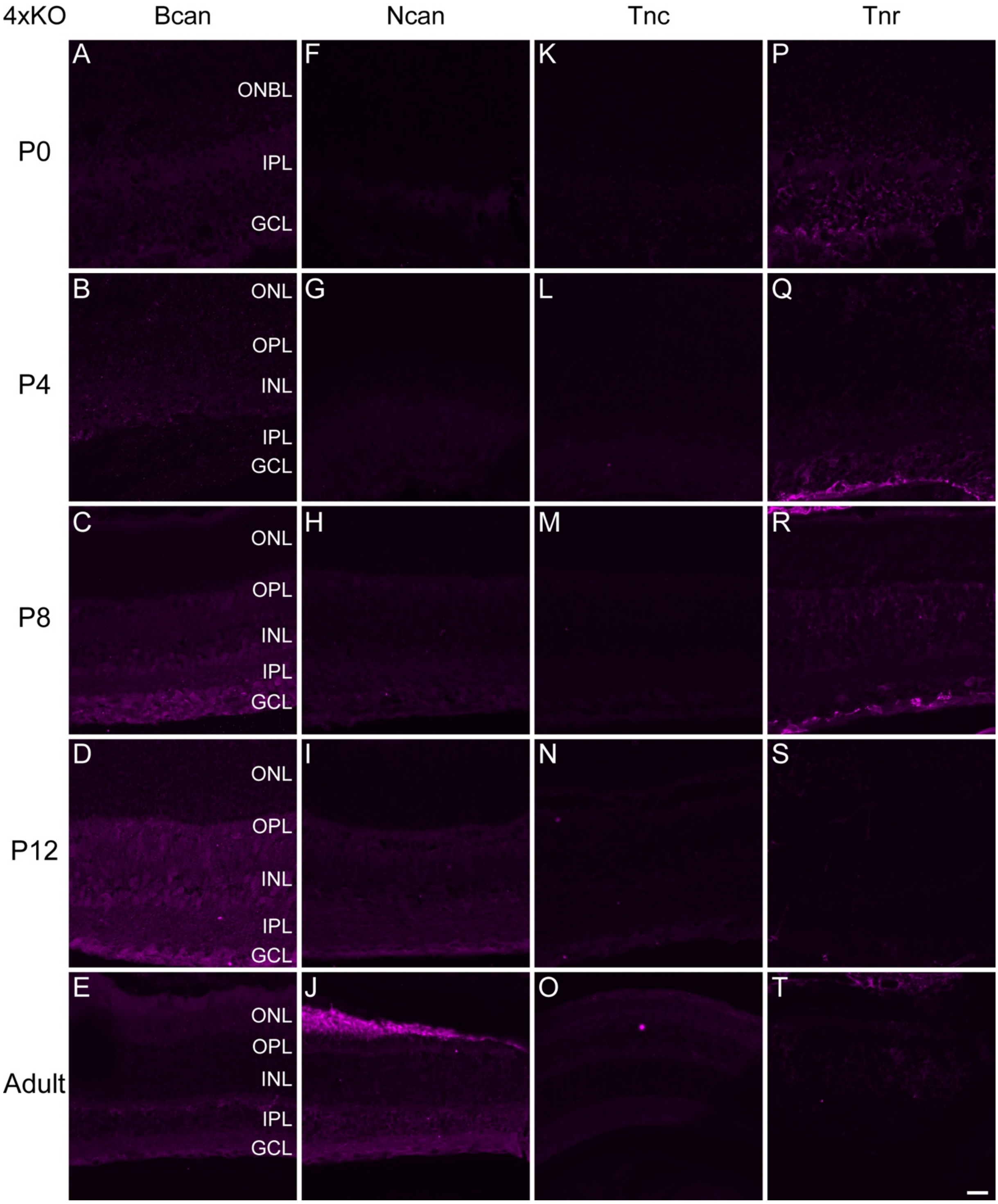


**Figure S2. Control staining of Bcan, Ncan, Tnc and Tnr in the postnatal and adult quadruple KO retina.**

Immunohistochemical staining of Bcan (A-E), Ncan (F-J), Tnc (K-O) and Tnr (P-T) in the postnatal (P0, P4, P8 and P12) and adult quadruple KO retina. The control staining revealed only faint, if any, background staining in the retina lacking the four molecules. Scale bar: 20 µm. 4xKO = quadruple knockout; Bcan = brevican; GCL = ganglion cell layer; INL = inner nuclear layer; IPL = inner plexiform layer; Ncan = neurocan; ONBL = outer neuroblastic layer; ONL = outer nuclear layer; OPL = outer plexiform layer; P = postnatal; Tnc = tenascin-C; Tnr = tenascin-R.

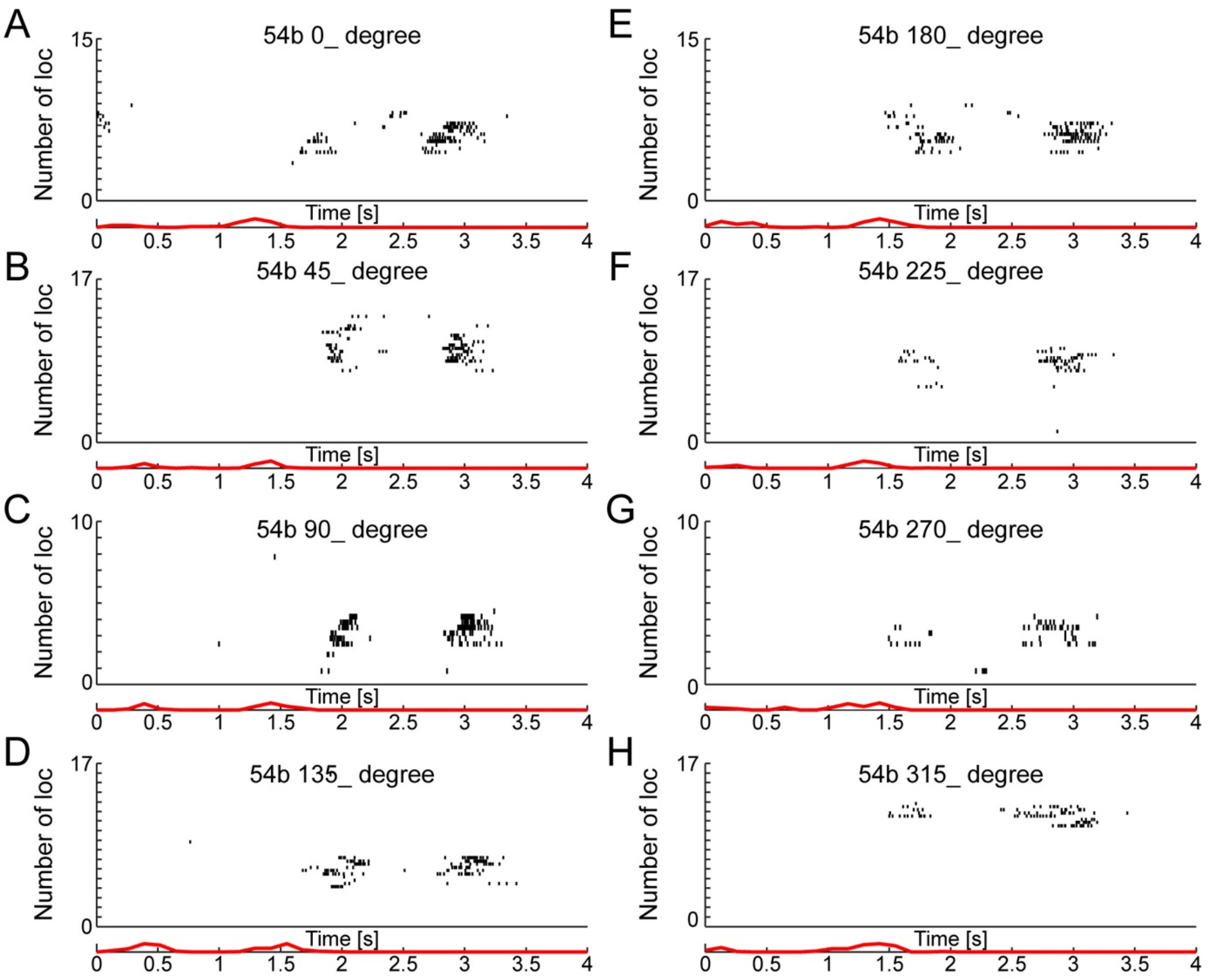


**Figure S3. Responses of an individual DSGC.** The diagrams depict the action potentials of an individual DSGC over time (s; 0°-315°). s = second.

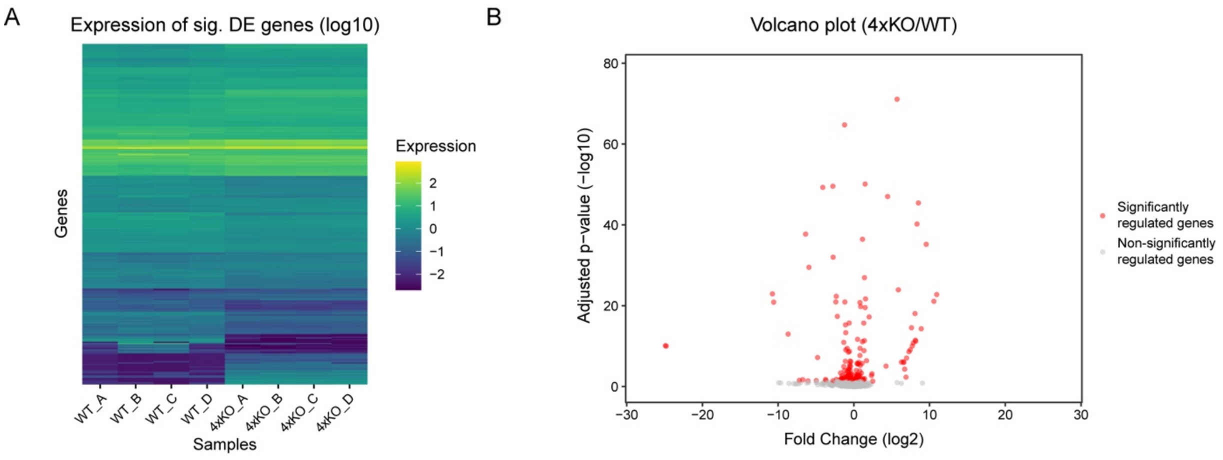


**Figure S4. NGS analyses identified 263 genes significantly altered in the quadruple KO compared to the WT retina.** (A) The heatmap illustrates the significant differentially 263 expressed genes (log10) in the WT and KO samples (N = 4). The expression levels of these genes are indicated by the color range from dark blue to yellow. 4xKO = quadruple knockout; WT = wildtype. (B) Additionally, a volcano plot shows significantly up- and downregulated genes (red dots) as well as non-significantly regulated genes (grey) according to their adjusted p-value (-log10) and fold change (log2).

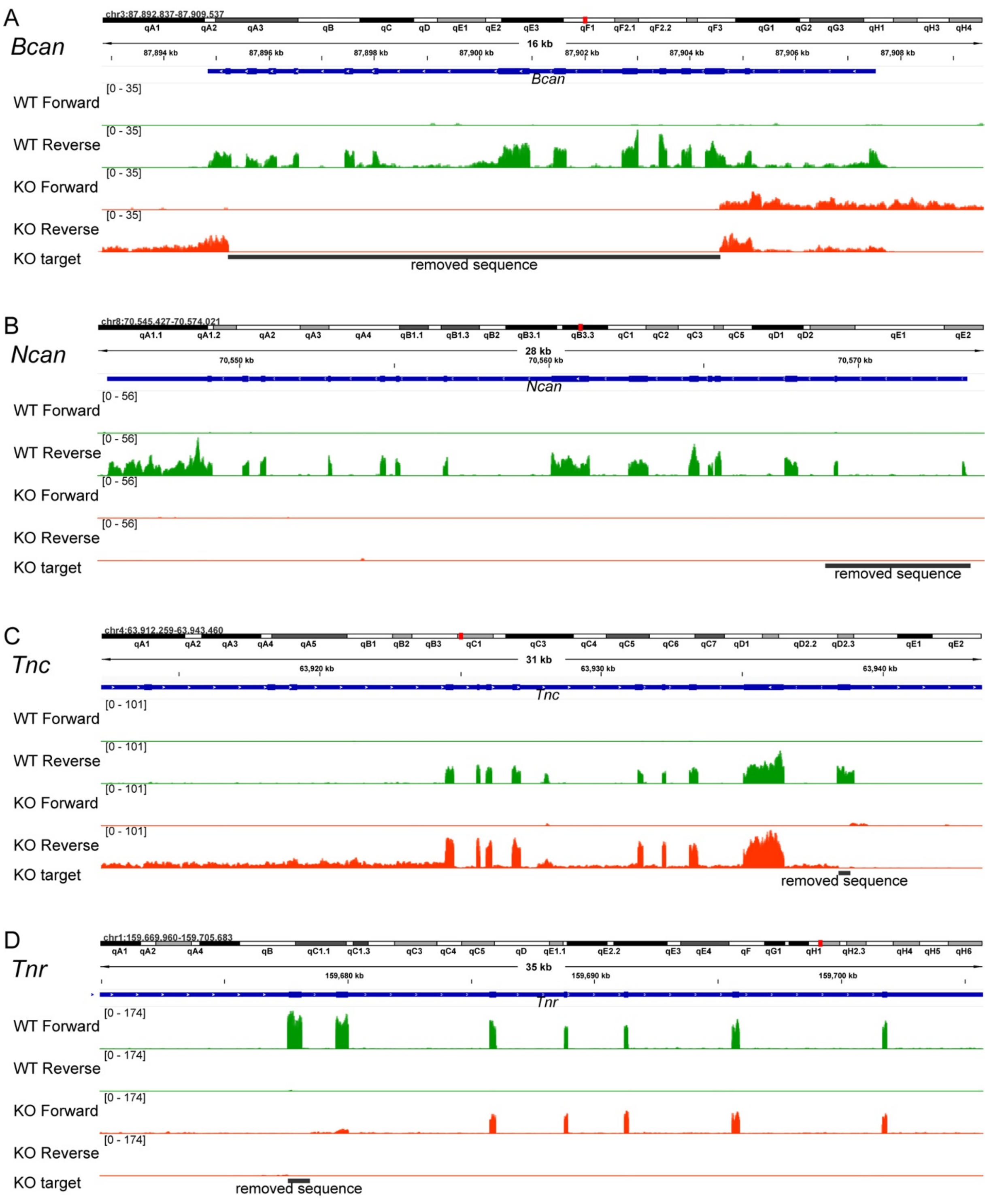


**Figure S5. Results of the Integrative Genomics Viewer (IGV) for the *Bcan, Ncan, Tnc* and *Tnr* alleles.** (A) The *Bcan* WT sequence is lost between position 87,895,223 and 87,904,572, according to GRCm39 (NC_000069.7). (B) The *Ncan* WT sequence is lost between position 70,568,941 and 70,573,644, according to GRCm39 (NC_000074.7). (C) The *Tnc* WT sequence is lost between position 63,938,412 and 63,938,833, according to GRCm39 (NC_000070.7). (D) The *Tnr* WT sequence is lost between position 159,677,592 and 159,678,477, according to GRCm39 (NC_000067.7). (A-D) The WT is shown in green, while the KO is shown in red. KO = knockout; WT = wildtype.

**Table S1. A- and b-wave amplitudes recorded from adult quadruple KO and WT mice *via* ERG analyses**. Light flash intensity, amplitude mean, SEM, genotypes and the p-value are indicated. P-values < 0.05 are shown in bold. 4xKO = quadruple knockout; WT = wildtype.

| **Light flash intensity [cd x s/m²]** | **0.1** | | **0.3** | | **1** | | **3** | | **10** | | **25** | |
|---|---|---|---|---|---|---|---|---|---|---|---|---|
| **Amplitude [µV]** | **Mean** | **SEM** | **Mean** | **SEM** | **Mean** | **SEM** | **Mean** | **SEM** | **Mean** | **SEM** | **Mean** | **SEM** |
| **A-wave** | | | | | | | | | | | | |
| WT | 75.87 | 6.41 | 133.04 | 7.00 | 155.64 | 8.00 | 162.14 | 10.25 | 186.08 | 10.25 | 202.57 | 9.75 |
| 4xKO | 59.36 | 3.88 | 107.09 | 5.82 | 129.61 | 6.78 | 144.28 | 5.47 | 156.36 | 8.73 | 184.60 | 11.03 |
| P-value | **0.035** | | **0.007** | | **0.018** | | **0.035** | | **0.035** | | **0.232** | |
| **B-wave** | | | | | | | | | | | | |
| WT | 483.34 | 19.64 | 552.33 | 26.26 | 560.28 | 25.80 | 517.33 | 21.68 | 618.24 | 29.80 | 628.26 | 29.11 |
| 4xKO | 341.19 | 17.95 | 404.38 | 17.50 | 425.27 | 19.20 | 408.29 | 18.03 | 464.41 | 19.81 | 487.53 | 16.91 |
| P-value | **< 0.001** | | **< 0.001** | | **< 0.001** | | **< 0.001** | | **< 0.001** | | **< 0.001** | |

### Table S2. Most prominently down- and upregulated genes (adjusted p-value < 0.05) in the retina of quadruple KO mice compared to WT mice as revealed by NGS.

| Ensembl ID | Gene symbol | Gene name | Fold change (log2) | Adjusted p-value |
|---|---|---|---|---|
| ENSMUSG00000119262 | *Snord3b1* | *Small nucleolar RNA, C/D box 3B1* | -24.899 | 7.64E-11 |
| ENSMUSG00000099329 | *NA* | *NA* | -24.778 | 9.56E-11 |
| ENSMUSG00000002341 | *Ncan* | *Neurocan* | -10.752 | 1.15E-23 |
| ENSMUSG00000117786 | *NA* | *NA* | -10.589 | 1.31E-21 |
| ENSMUSG00000020062 | *Slc5a8* | *Solute carrier family 5 (iodide transporter), member 8* | -8.683 | 1.02E-13 |
| ENSMUSG00000116347 | *NA* | *NA* | -7.202 | 0.0290256 |
| ENSMUSG00000058625 | *Gm17383* | *Tubulin, alpha 1B pseudogene* | -6.748 | 0.0202855 |
| ENSMUSG00000114982 | *Vmn1r14* | *Vomeronasal 1 receptor 14* | -6.374 | 1.86E-38 |
| ENSMUSG00000081067 | *Gm12912* | *Eukaryotic translation elongation factor 1 alpha 1 pseudogene* | -6.063 | 0.03902883 |
| ENSMUSG00000024027 | *Glp1r* | *Glucagon-like peptide 1 receptor* | -5.933 | 3.08E-30 |
| ENSMUSG00000083311 | *NA* | *NA* | -4.964 | 0.03243631 |
| ENSMUSG00000115199 | *Vmn1r15* | *Vomeronasal 1 receptor 15* | -4.791 | 6.50E-08 |
| ENSMUSG00000004892 | *Bcan* | *Brevican* | -4.103 | 5.10E-50 |
| ENSMUSG00000117468 | *Gm7059* | *Predicted gene 7059* | -3.740 | 0.01751706 |
| ENSMUSG00000074978 | *Actg-ps1* | *Actin, gamma, pseudogene 1* | -3.700 | 0.03398373 |
| ENSMUSG00000004070 | *Hmox2* | *Heme oxygenase 2* | -2.778 | 2.68E-50 |
| ENSMUSG00000118193 | *NA* | *NA* | -2.744 | 0.03533261 |
| ENSMUSG00000104988 | *NA* | *NA* | -2.741 | 9.73E-33 |
| ENSMUSG00000033826 | *Dnah8* | *Dynein, axonemal, heavy chain 8* | -2.382 | 1.15E-21 |
| ENSMUSG00000001930 | *Vwf* | *Von Willebrand factor* | -2.329 | 4.73E-23 |
| ENSMUSG00000088025 | *Rprl3* | *Ribonuclease P RNA-like 3* | -2.307 | 0.01667936 |
| ENSMUSG00000001520 | *Nrip2* | *Nuclear receptor interacting protein 2* | -2.183 | 4.23E-18 |
| ENSMUSG00000027570 | *Col9a3* | *Collagen, type IX, alpha 3* | -2.075 | 0.03386148 |
| ENSMUSG00000092805 | *NA* | *NA* | -1.804 | 0.03386148 |
| ENSMUSG00000081400 | *Gm13680* | *Ribosomal protein S4, X-linked pseudogene* | -1.767 | 0.04109815 |
| ENSMUSG00000117430 | *NA* | *NA* | -1.759 | 0.00042564 |
| ENSMUSG00000035238 | *Kcnk15* | *Potassium channel, subfamily K, member 15* | -1.686 | 0.04294251 |
| ENSMUSG00000029468 | *P2rx7* | *Purinergic receptor P2X, ligand-gated ion channel, 7* | -1.683 | 0.01104677 |
| ENSMUSG00000100690 | *NA* | *NA* | -1.672 | 0.0481746 |
| ENSMUSG00000085042 | *Abhd11os* | *Abhydrolase domain containing 11, opposite strand* | -1.670 | 0.01816235 |
| ENSMUSG00000099041 | *NA* | *NA* | -1.622 | 0.01419505 |
| ENSMUSG00000113123 | *NA* | *NA* | -1.580 | 0.00811916 |
| ENSMUSG00000099696 | *2900052N01Rik* | *RIKEN cdna 2900052N01 gene* | -1.541 | 0.00195106 |
| ENSMUSG00000112759 | *NA* | *NA* | -1.527 | 0.03384861 |
| ENSMUSG00000028339 | *Col15a1* | *Collagen, type XV, alpha 1* | -1.462 | 0.03736654 |
| ENSMUSG00000107023 | *NA* | *NA* | -1.436 | 0.03074642 |
| ENSMUSG00000051747 | *Ttn* | *Titin* | -1.425 | 0.02321926 |

| | | | | |
|---|---|---|---|---|
| ENSMUSG00000040957 | *Cables1* | *CDK5 and Abl enzyme substrate 1* | -1.337 | 1.17E-11 |
| ENSMUSG00000034762 | *Glis1* | *GLIS family zinc finger 1* | -1.328 | 1.09E-05 |
| ENSMUSG00000037375 | *Hhat* | *Hedgehog acyltransferase* | -1.309 | 0.00976939 |
| ENSMUSG00000024330 | *Col11a2* | *Collagen, type XI, alpha 2* | -1.290 | 0.00595575 |
| ENSMUSG00000071341 | *Egr4* | *Early growth response 4* | -1.282 | 0.01049129 |
| ENSMUSG00000065176 | *Rnu12* | *RNA U12, small nuclear* | -1.272 | 0.03602475 |
| ENSMUSG00000078496 | *Zfp982* | *Zinc finger protein 982* | -1.260 | 0.03077515 |
| ENSMUSG00000035606 | *Ky* | *Kyphoscoliosis peptidase* | -1.248 | 0.0001027 |
| ENSMUSG00000015829 | *Tnr* | *Tenascin R* | -1.223 | 1.76E-65 |
| ENSMUSG00000024026 | *Glo1* | *Glyoxalase 1* | -1.189 | 1.17E-21 |
| ENSMUSG00000089417 | *Gm22009* | *Predicted gene, 22009* | -1.152 | 0.04087902 |
| ENSMUSG00000004891 | *Nes* | *Nestin* | -1.124 | 5.40E-07 |
| ENSMUSG00000117725 | *NA* | *NA* | -1.103 | 5.35E-16 |
| ENSMUSG00000019935 | *Slc17a8* | *Solute carrier family 17 (sodium-dependent inorganic phosphate cotransporter), member 8* | -1.069 | 8.33E-10 |
| ENSMUSG00000097440 | *NA* | *NA* | -1.068 | 4.57E-14 |
| ENSMUSG00000044005 | *Gls2* | *Glutaminase 2 (liver, mitochondrial)* | -1.068 | 0.00065531 |
| ENSMUSG00000056724 | *Nbeal2* | *Neurobeachin-like 2* | -1.029 | 0.03376583 |
| ENSMUSG00000098087 | *Gm17750* | *Predicted gene, 17750* | -1.012 | 0.03398373 |
| ENSMUSG00000017929 | *B4galt5* | *UDP-Gal:betaglcnac beta 1,4-galactosyltransferase, polypeptide 5* | -1.009 | 0.02960603 |
| ENSMUSG00000111758 | *NA* | *NA* | -1.005 | 0.01874757 |
| ENSMUSG00000092837 | *Rpph1* | *Ribonuclease P RNA component H1* | -0.975 | 0.03097316 |
| ENSMUSG00000026042 | *Col5a2* | *Collagen, type V, alpha 2* | -0.902 | 0.01098075 |
| ENSMUSG00000024378 | *Stard4* | *Star-related lipid transfer (START) domain containing 4* | -0.878 | 3.53E-10 |
| ENSMUSG00000021420 | *Fars2* | *Phenylalanine-trna synthetase 2 (mitochondrial)* | -0.864 | 0.00203584 |
| ENSMUSG00000007594 | *Hapln4* | *Hyaluronan and proteoglycan link protein 4* | -0.864 | 0.00338482 |
| ENSMUSG00000051435 | *Fhad1* | *Forkhead-associated (FHA) phosphopeptide binding domain 1* | -0.855 | 0.03913768 |
| ENSMUSG00000041258 | *Zfp236* | *Zinc finger protein 236* | -0.852 | 0.03193908 |
| ENSMUSG00000018849 | *Wwc1* | *WW, C2 and coiled-coil domain containing 1* | -0.836 | 0.00012095 |
| ENSMUSG00000044471 | *Lncpint* | *Long non-protein coding RNA. Trp53 induced transcript* | -0.820 | 0.008272 |
| ENSMUSG00000032558 | *Nphp3* | *Nephronophthisis 3 (adolescent)* | -0.820 | 0.03386148 |
| ENSMUSG00000056899 | *Immp2l* | *IMP2 inner mitochondrial membrane peptidase-like (S. Cerevisiae)* | -0.819 | 0.0392063 |
| ENSMUSG00000097412 | *1810014B01Rik* | *RIKEN cdna 1810014B01 gene* | -0.817 | 0.0069357 |
| ENSMUSG00000037010 | *Apln* | *Apelin* | -0.766 | 0.00620343 |
| ENSMUSG00000026604 | *Ptpn14* | *Protein tyrosine phosphatase, non-receptor type 14* | -0.760 | 0.00196781 |
| ENSMUSG00000024421 | *Lama3* | *Laminin, alpha 3* | -0.748 | 0.0392063 |
| ENSMUSG00000046402 | *Rbp1* | *Retinol binding protein 1, cellular* | -0.745 | 0.03814001 |
| ENSMUSG00000019899 | *Lama2* | *Laminin, alpha 2* | -0.742 | 0.02917959 |
| ENSMUSG00000071112 | *Spx* | *Spexin hormone* | -0.732 | 0.00226575 |
| ENSMUSG00000062202 | *Btbd9* | *BTB (POZ) domain containing 9* | -0.722 | 3.05E-09 |
| ENSMUSG00000006281 | *Tep1* | *Telomerase associated protein 1* | -0.719 | 0.02632381 |
| ENSMUSG00000022129 | *Dct* | *Dopachrome tautomerase* | -0.716 | 0.00016851 |
| ENSMUSG00000026098 | *Pms1* | *PMS1 homolog 1, mismatch repair system component* | -0.689 | 0.03376583 |

| | | | | |
|---|---|---|---|---|
| ENSMUSG00000061533 | *Cep128* | *Centrosomal protein 128* | -0.678 | 0.03043674 |
| ENSMUSG00000023723 | *Mrps23* | *Mitochondrial ribosomal protein S23* | -0.672 | 0.02633536 |
| ENSMUSG00000009076 | *Zmat5* | *Zinc finger, matrin type 5* | -0.668 | 4.37E-05 |
| ENSMUSG00000052921 | *Arhgef15* | *Rho guanine nucleotide exchange factor (GEF) 15* | -0.666 | 0.04237172 |
| ENSMUSG00000024404 | *Riok3* | *RIO kinase 3* | -0.666 | 0.03757008 |
| ENSMUSG00000015980 | *Lrrc27* | *Leucine rich repeat containing 27* | -0.664 | 5.65E-05 |
| ENSMUSG00000028391 | *Wdr31* | *WD repeat domain 31* | -0.664 | 0.01524945 |
| ENSMUSG00000024713 | *Pcsk5* | *Proprotein convertase subtilisin/kexin type 5* | -0.637 | 0.00138565 |
| ENSMUSG00000090799 | *Klhl33* | *Kelch-like 33* | -0.632 | 0.01741592 |
| ENSMUSG00000048826 | *Dact2* | *Dishevelled-binding antagonist of beta-catenin 2* | -0.624 | 0.04351747 |
| ENSMUSG00000078566 | *Bnip3* | *BCL2/adenovirus E1B interacting protein 3* | -0.617 | 1.88E-16 |
| ENSMUSG00000043822 | *Adamtsl5* | *ADAMTS-like 5* | -0.613 | 0.03610635 |
| ENSMUSG00000033209 | *Ttc28* | *Tetratricopeptide repeat domain 28* | -0.604 | 0.00496804 |
| ENSMUSG00000061911 | *Myt1l* | *Myelin transcription factor 1-like* | -0.602 | 0.04036859 |
| ENSMUSG00000039410 | *Prdm16* | *PR domain containing 16* | -0.598 | 0.02216696 |
| ENSMUSG00000030230 | *Plcz1* | *Phospholipase C, zeta 1* | -0.593 | 0.00165668 |
| ENSMUSG00000024228 | *Nudt12* | *Nudix (nucleoside diphosphate linked moiety X)-type motif 12* | -0.584 | 0.03043674 |
| ENSMUSG00000041957 | *Pkp2* | *Plakophilin 2* | -0.581 | 2.01E-09 |
| ENSMUSG00000014846 | *Tppp3* | *Tubulin polymerization-promoting protein family member 3* | -0.576 | 0.04126425 |
| ENSMUSG00000024287 | *Thoc1* | *THO complex 1* | -0.572 | 0.00184145 |
| ENSMUSG00000027070 | *Lrp2* | *Low density lipoprotein receptor-related protein 2* | -0.564 | 0.00078386 |
| ENSMUSG00000040811 | *Eml2* | *Echinoderm microtubule associated protein like 2* | -0.562 | 0.04294251 |
| ENSMUSG00000097391 | *Mirg* | *Mirna containing gene* | -0.551 | 0.01935485 |
| ENSMUSG00000000628 | *Hk2* | *Hexokinase 2* | -0.549 | 9.86E-06 |
| ENSMUSG00000054477 | *Kcnn2* | *Potassium intermediate/small conductance calcium-activated channel, subfamily N, member 2* | -0.548 | 2.86E-06 |
| ENSMUSG00000032387 | *Rbpms2* | *RNA binding protein with multiple splicing 2* | -0.523 | 0.01972036 |
| ENSMUSG00000024942 | *Capn1* | *Calpain 1* | -0.516 | 0.02775487 |
| ENSMUSG00000038539 | *Atf5* | *Activating transcription factor 5* | -0.510 | 0.01184271 |
| ENSMUSG00000002602 | *Axl* | *AXL receptor tyrosine kinase* | -0.502 | 0.03656956 |
| ENSMUSG00000036879 | *Phkb* | *Phosphorylase kinase beta* | -0.499 | 0.02596979 |
| ENSMUSG00000020926 | *Adam11* | *A disintegrin and metallopeptidase domain 11* | -0.497 | 0.03097316 |
| ENSMUSG00000024413 | *Npc1* | *NPC intracellular cholesterol transporter 1* | -0.490 | 4.84E-07 |
| ENSMUSG00000118667 | *Ahnak2* | *AHNAK nucleoprotein 2* | -0.483 | 0.03386148 |
| ENSMUSG00000044982 | *Sft2d3* | *SFT2 domain containing 3* | -0.480 | 0.03707452 |
| ENSMUSG00000016346 | *Kcnq2* | *Potassium voltage-gated channel, subfamily Q, member 2* | -0.479 | 0.0481746 |
| ENSMUSG00000029510 | *Gpc2* | *Glypican 2 (cerebroglycan)* | -0.477 | 0.04715658 |
| ENSMUSG00000049044 | *Rapgef4* | *Rap guanine nucleotide exchange factor (GEF) 4* | -0.477 | 8.85E-05 |
| ENSMUSG00000113216 | *NA* | *NA* | -0.471 | 0.02906688 |
| ENSMUSG00000024293 | *Esco1* | *Establishment of sister chromatid cohesion N-acetyltransferase 1* | -0.449 | 9.04E-06 |
| ENSMUSG00000040249 | *Lrp1* | *Low density lipoprotein receptor-related protein 1* | -0.446 | 0.0481746 |
| ENSMUSG00000042515 | *Pwwp3b* | *PWWP domain containing 3B* | -0.446 | 0.03821277 |
| ENSMUSG00000024294 | *Mib1* | *Mindbomb E3 ubiquitin protein ligase 1* | -0.437 | 0.01574217 |

| | | | | |
|---|---|---|---|---|
| ENSMUSG00000031586 | *Rbpms* | *RNA binding protein gene with multiple splicing* | -0.426 | 0.03898348 |
| ENSMUSG00000051401 | *Kctd16* | *Potassium channel tetramerisation domain containing 16* | -0.422 | 0.04381177 |
| ENSMUSG00000047879 | *Usp14* | *Ubiquitin specific peptidase 14* | -0.413 | 6.96E-07 |
| ENSMUSG00000055471 | *Alk* | *Anaplastic lymphoma kinase* | -0.397 | 0.01891562 |
| ENSMUSG00000019775 | *Rgs17* | *Regulator of G-protein signaling 17* | -0.389 | 0.00333549 |
| ENSMUSG00000028937 | *Acot7* | *Acyl-coa thioesterase 7* | -0.380 | 0.03097316 |
| ENSMUSG00000024082 | *Ndufaf7* | *NADH:ubiquinone oxidoreductase complex assembly factor 7* | -0.370 | 0.02028546 |
| ENSMUSG00000029001 | *Fbxo44* | *F-box protein 44* | -0.350 | 0.04647076 |
| ENSMUSG00000071337 | *Tia1* | *Cytotoxic granule-associated RNA binding protein 1* | -0.347 | 0.04087902 |
| ENSMUSG00000048148 | *Nwd1* | *NACHT and WD repeat domain containing 1* | -0.345 | 0.03386148 |
| ENSMUSG00000020305 | *Asb3* | *Ankyrin repeat and SOCS box-containing 3* | -0.343 | 0.02090752 |
| ENSMUSG00000011751 | *Sptbn4* | *Spectrin beta non-erythrocytic 4* | -0.332 | 0.04194938 |
| ENSMUSG00000005533 | *Igf1r* | *Insulin-like growth factor I receptor* | -0.326 | 0.00988524 |
| ENSMUSG00000002475 | *Abhd3* | *Abhydrolase domain containing 3* | -0.322 | 0.03382175 |
| ENSMUSG00000019828 | *Grm1* | *Glutamate receptor, metabotropic 1* | -0.317 | 0.0010475 |
| ENSMUSG00000006740 | *Kif5b* | *Kinesin family member 5B* | -0.289 | 0.03814001 |
| ENSMUSG00000022043 | *Trim35* | *Tripartite motif-containing 35* | -0.284 | 0.0392063 |
| ENSMUSG00000079509 | *Zfx* | *Zinc finger protein X-linked* | -0.269 | 0.04303407 |
| ENSMUSG00000051510 | *Mafg* | *V-maf musculoaponeurotic fibrosarcoma oncogene family, protein G (avian)* | -0.269 | 0.008272 |
| ENSMUSG00000056268 | *Dennd1b* | *DENN/MADD domain containing 1B* | -0.268 | 0.00078386 |
| ENSMUSG00000021196 | *Pfkp* | *Phosphofructokinase, platelet* | -0.182 | 0.01390382 |
| ENSMUSG00000036371 | *Serbp1* | *Serpine1 mrna binding protein 1* | 0.148 | 0.03290012 |
| ENSMUSG00000029328 | *Hnrnpdl* | *Heterogeneous nuclear ribonucleoprotein D-like* | 0.166 | 0.03814001 |
| ENSMUSG00000068747 | *Sort1* | *Sortilin 1* | 0.202 | 0.03193908 |
| ENSMUSG00000060733 | *Ipmk* | *Inositol polyphosphate multikinase* | 0.236 | 0.03757008 |
| ENSMUSG00000096351 | *Samd11* | *Sterile alpha motif domain containing 11* | 0.242 | 0.01651863 |
| ENSMUSG00000027634 | *Ndrg3* | *N-myc downstream regulated gene 3* | 0.261 | 0.00888043 |
| ENSMUSG00000009905 | *Kdsr* | *3-ketodihydrosphingosine reductase* | 0.261 | 0.0392063 |
| ENSMUSG00000020700 | *Map3k3* | *Mitogen-activated protein kinase kinase kinase 3* | 0.263 | 0.01474962 |
| ENSMUSG00000035266 | *Helq* | *Helicase, POLQ-like* | 0.267 | 0.04193705 |
| ENSMUSG00000046593 | *Tmem215* | *Transmembrane protein 215* | 0.267 | 0.0263148 |
| ENSMUSG00000029410 | *Ppef2* | *Protein phosphatase, EF hand calcium-binding domain 2* | 0.281 | 0.01049129 |
| ENSMUSG00000048222 | *Mfap1b* | *Microfibrillar-associated protein 1B* | 0.288 | 0.02458349 |
| ENSMUSG00000066735 | *Vkorc1l1* | *Vitamin K epoxide reductase complex, subunit 1-like 1* | 0.289 | 0.00779139 |
| ENSMUSG00000003929 | *Zfp81* | *Zinc finger protein 81* | 0.295 | 0.03814001 |
| ENSMUSG00000021803 | *Cdhr1* | *Cadherin-related family member 1* | 0.303 | 0.04417455 |
| ENSMUSG00000008301 | *Phax* | *Phosphorylated adaptor for RNA export* | 0.303 | 0.03386148 |
| ENSMUSG00000058966 | *Tlcd3b* | *TLC domain containing 3B* | 0.304 | 0.03376583 |
| ENSMUSG00000022332 | *Khdrbs3* | *KH domain containing, RNA binding, signal transduction associated 3* | 0.306 | 0.03393176 |
| ENSMUSG00000036779 | *Tent4b* | *Terminal nucleotidyltransferase 4B* | 0.309 | 0.04294251 |
| ENSMUSG00000054893 | *Zfp667* | *Zinc finger protein 667* | 0.314 | 0.04792744 |
| ENSMUSG00000029513 | *Prkab1* | *Protein kinase, AMP-activated, beta 1 non-catalytic subunit* | 0.318 | 0.0001027 |

| | | | | |
|---|---|---|---|---|
| ENSMUSG00000024824 | *Rad9a* | *RAD9 checkpoint clamp component A* | 0.319 | 0.02028546 |
| ENSMUSG00000022131 | *Gpr180* | *G protein-coupled receptor 180* | 0.321 | 0.03611457 |
| ENSMUSG00000032501 | *Trib1* | *Tribbles pseudokinase 1* | 0.325 | 0.02321926 |
| ENSMUSG00000008035 | *Mid1ip1* | *Mid1 interacting protein 1 (gastrulation specific G12-like (zebrafish))* | 0.339 | 0.03942002 |
| ENSMUSG00000021189 | *Atxn3* | *Ataxin 3* | 0.341 | 0.00398635 |
| ENSMUSG00000051502 | *Ufsp1* | *UFM1-specific peptidase 1* | 0.344 | 0.03193908 |
| ENSMUSG00000020034 | *Tcp11l2* | *T-complex 11 (mouse) like 2* | 0.360 | 0.00363053 |
| ENSMUSG00000025277 | *Abhd6* | *Abhydrolase domain containing 6* | 0.367 | 0.01476881 |
| ENSMUSG00000026618 | *Iars2* | *Isoleucine-trna synthetase 2, mitochondrial* | 0.368 | 0.01564914 |
| ENSMUSG00000004655 | *Aqp1* | *Aquaporin 1* | 0.370 | 0.00479062 |
| ENSMUSG00000053773 | *Rdh8* | *Retinol dehydrogenase 8* | 0.372 | 0.03942002 |
| ENSMUSG00000025380 | *Fscn2* | *Fascin actin-bundling protein 2* | 0.377 | 0.02823471 |
| ENSMUSG00000103332 | *Pcdhga2* | *Protocadherin gamma subfamily A, 2* | 0.385 | 0.01524945 |
| ENSMUSG00000024238 | *Zeb1* | *Zinc finger E-box binding homeobox 1* | 0.395 | 0.00099531 |
| ENSMUSG00000024900 | *Cpt1a* | *Carnitine palmitoyltransferase 1a, liver* | 0.397 | 0.02964575 |
| ENSMUSG00000035504 | *Reep6* | *Receptor accessory protein 6* | 0.412 | 0.00241542 |
| ENSMUSG00000031540 | *Kat6a* | *K(lysine) acetyltransferase 6A* | 0.412 | 0.03814001 |
| ENSMUSG00000033396 | *Spg11* | *SPG11, spatacsin vesicle trafficking associated* | 0.413 | 1.80E-06 |
| ENSMUSG00000041225 | *Arhgap12* | *Rho gtpase activating protein 12* | 0.419 | 0.00109718 |
| ENSMUSG00000036570 | *Fxyd1* | *FXYD domain-containing ion transport regulator 1* | 0.419 | 0.04294251 |
| ENSMUSG00000092416 | *Zfp141* | *Zinc finger protein 141* | 0.424 | 0.03386355 |
| ENSMUSG00000022724 | *Riox2* | *Ribosomal oxygenase 2* | 0.424 | 0.02134901 |
| ENSMUSG00000034450 | *Gulo* | *Gulonolactone (L-) oxidase* | 0.434 | 0.02775487 |
| ENSMUSG00000074037 | *Mc1r* | *Melanocortin 1 receptor* | 0.444 | 0.04299375 |
| ENSMUSG00000043639 | *Rbm20* | *RNA binding motif protein 20* | 0.448 | 0.00269897 |
| ENSMUSG00000021203 | *Otub2* | *OTU domain, ubiquitin aldehyde binding 2* | 0.449 | 0.01599243 |
| ENSMUSG00000070368 | *Prok1* | *Prokineticin 1* | 0.449 | 0.02301935 |
| ENSMUSG00000035620 | *Ric8b* | *RIC8 guanine nucleotide exchange factor B* | 0.457 | 0.04193705 |
| ENSMUSG00000100252 | *Mir124-2hg* | *Mir124-2 host gene (non-protein coding)* | 0.462 | 0.04779972 |
| ENSMUSG00000028328 | *Tmod1* | *Tropomodulin 1* | 0.468 | 0.01343914 |
| ENSMUSG00000024206 | *Rfx2* | *Regulatory factor X, 2 (influences HLA class II expression)* | 0.480 | 0.02090752 |
| ENSMUSG00000047298 | *Kcnv2* | *Potassium channel, subfamily V, member 2* | 0.482 | 0.04417455 |
| ENSMUSG00000074794 | *Arrdc3* | *Arrestin domain containing 3* | 0.485 | 0.00184145 |
| ENSMUSG00000038141 | *Tmem181a* | *Transmembrane protein 181A* | 0.486 | 2.06E-12 |
| ENSMUSG00000035769 | *Xylb* | *Xylulokinase homolog (H. Influenzae)* | 0.493 | 0.03598918 |
| ENSMUSG00000054237 | *Fra10ac1* | *FRA10AC1 homolog (human)* | 0.496 | 0.02028546 |
| ENSMUSG00000112649 | *NA* | *NA* | 0.502 | 0.03814001 |
| ENSMUSG00000034453 | *Polr3b* | *Polymerase (RNA) III (DNA directed) polypeptide B* | 0.514 | 2.64E-06 |
| ENSMUSG00000074829 | *2010315B03Rik* | *RIKEN cdna 2010315B03 gene* | 0.535 | 0.04972459 |
| ENSMUSG00000048416 | *Mlf1* | *Myeloid leukemia factor 1* | 0.577 | 2.64E-06 |
| ENSMUSG00000028629 | *Exo5* | *Exonuclease 5* | 0.617 | 0.01512724 |
| ENSMUSG00000039037 | *St6galnac5* | *ST6(alpha-N-acetyl-neuraminyl-2,3-beta-galactosyl-1,3)-N-acetylgalactosaminide alpha-2,6-sialyltransferase 5* | 0.640 | 0.02964047 |

| | | | | |
|---|---|---|---|---|
| ENSMUSG00000041891 | *Lman1* | *Lectin, mannose-binding, 1* | 0.641 | 9.96E-07 |
| ENSMUSG00000026154 | *Sdhaf4* | *Succinate dehydrogenase complex assembly factor 4* | 0.656 | 0.00226575 |
| ENSMUSG00000082286 | *NA* | *NA* | 0.659 | 0.00233415 |
| ENSMUSG00000018263 | *Tbx5* | *T-box 5* | 0.661 | 0.03097316 |
| ENSMUSG00000041238 | *Rbbp8* | *Retinoblastoma binding protein 8, endonuclease* | 0.766 | 0.02906822 |
| ENSMUSG00000000579 | *Dynlt1c* | *Dynein light chain Tctex-type 1C* | 0.780 | 0.00014602 |
| ENSMUSG00000032468 | *Armc8* | *Armadillo repeat containing 8* | 0.800 | 2.02E-08 |
| ENSMUSG00000043773 | *1700048O20Rik* | *RIKEN cdna 1700048O20 gene* | 0.801 | 0.00176454 |
| ENSMUSG00000026156 | *B3gat2* | *Beta-1,3-glucuronyltransferase 2 (glucuronosyltransferase S)* | 0.802 | 1.62E-21 |
| ENSMUSG00000026141 | *Col19a1* | *Collagen, type XIX, alpha 1* | 0.814 | 3.53E-10 |
| ENSMUSG00000086629 | *2810403D21Rik* | *RIKEN cdna 2810403D21 gene* | 0.843 | 0.00976939 |
| ENSMUSG00000024260 | *Sap130* | *Sin3A associated protein* | 0.853 | 5.63E-06 |
| ENSMUSG00000032527 | *Pccb* | *Propionyl Coenzyme A carboxylase, beta polypeptide* | 0.899 | 1.58E-20 |
| ENSMUSG00000033581 | *Igf2bp2* | *Insulin-like growth factor 2 mrna binding protein 2* | 0.995 | 0.00047722 |
| ENSMUSG00000093880 | *NA* | *NA* | 1.080 | 1.78E-06 |
| ENSMUSG00000092074 | *Dynlt1a* | *Dynein light chain Tctex-type 1A* | 1.135 | 7.92E-12 |
| ENSMUSG00000026155 | *Smap1* | *Small arfgap 1* | 1.138 | 3.54E-37 |
| ENSMUSG00000084838 | *NA* | *NA* | 1.151 | 0.0404933 |
| ENSMUSG00000038541 | *Srd5a2* | *Steroid 5 alpha-reductase 2* | 1.222 | 0.0117113 |
| ENSMUSG00000038094 | *Atp13a4* | *Atpase type 13A4* | 1.239 | 0.02650919 |
| ENSMUSG00000109176 | *NA* | *NA* | 1.242 | 0.02090752 |
| ENSMUSG00000026153 | *Fam135a* | *Family with sequence similarity 135, member A* | 1.307 | 1.96E-16 |
| ENSMUSG00000026158 | *Ogfrl1* | *Opioid growth factor receptor-like 1* | 1.341 | 1.30E-09 |
| ENSMUSG00000024992 | *Pde6c* | *Phosphodiesterase 6C, cgmp specific, cone, alpha prime* | 1.390 | 4.31E-12 |
| ENSMUSG00000078886 | *Gm2026* | *Predicted gene 2026* | 1.397 | 1.12E-27 |
| ENSMUSG00000097387 | *4930563E18Rik* | *RIKEN cdna 4930563E18 gene* | 1.427 | 0.01126644 |
| ENSMUSG00000078899 | *Gm14430* | *Predicted gene 14430* | 1.487 | 7.60E-51 |
| ENSMUSG00000096780 | *NA* | *NA* | 1.491 | 2.78E-20 |
| ENSMUSG00000063953 | *Amd2* | *S-adenosylmethionine decarboxylase 2* | 1.537 | 2.02E-22 |
| ENSMUSG00000078897 | *Gm4724* | *Predicted gene 4724* | 1.672 | 4.00E-07 |
| ENSMUSG00000096255 | *Dynlt1b* | *Dynein light chain Tctex-type 1B* | 2.019 | 5.77E-18 |
| ENSMUSG00000078880 | *Gm14308* | *Predicted gene 14308* | 2.328 | 0.00195106 |
| ENSMUSG00000117494 | *NA* | *NA* | 2.390 | 0.00067931 |
| ENSMUSG00000095295 | *Gm3488* | *Predicted gene, 3488* | 2.489 | 0.04715658 |
| ENSMUSG00000115160 | *NA* | *NA* | 4.247 | 8.77E-06 |
| ENSMUSG00000078870 | *Gm14410* | *Predicted gene 14410* | 4.458 | 9.52E-48 |
| ENSMUSG00000072594 | *NA* | *NA* | 5.709 | 7.98E-72 |
| ENSMUSG00000078866 | *Zfp970* | *Zinc finger protein 970* | 5.875 | 1.15E-24 |
| ENSMUSG00000078875 | *NA* | *NA* | 6.283 | 9.08E-07 |
| ENSMUSG00000082765 | *NA* | *NA* | 6.532 | 7.43E-07 |
| ENSMUSG00000084858 | *NA* | *NA* | 6.604 | 1.12E-06 |
| ENSMUSG00000078894 | *2210418O10Rik* | *RIKEN cdna 2210418O10 gene* | 6.708 | 4.91E-05 |

| | | | | |
|---|---|---|---|---|
| ENSMUSG00000078896 | *Zfp965* | *Zinc finger protein 965* | 6.883 | 0.00469196 |
| ENSMUSG00000082724 | *NA* | *NA* | 6.962 | 8.56E-08 |
| ENSMUSG00000074521 | *NA* | *NA* | 7.275 | 2.37E-09 |
| ENSMUSG00000095648 | *Gm2004* | *Predicted gene 2004* | 7.450 | 8.33E-10 |
| ENSMUSG00000078879 | *Zfp973* | *Zinc finger protein 973* | 7.610 | 2.78E-15 |
| ENSMUSG00000094786 | *NA* | *NA* | 7.718 | 9.24E-11 |
| ENSMUSG00000083111 | *NA* | *NA* | 7.854 | 1.62E-11 |
| ENSMUSG00000074519 | *Zfp971* | *Zinc finger protein 971* | 8.076 | 8.57E-19 |
| ENSMUSG00000078868 | *Gm14412* | *Predicted gene 14412* | 8.117 | 3.39E-12 |
| ENSMUSG00000078864 | *Gm14322* | *Predicted gene 14322* | 8.173 | 6.18E-12 |
| ENSMUSG00000078878 | *Gm14305* | *Predicted gene 14305* | 8.339 | 6.04E-41 |
| ENSMUSG00000074527 | *Gm14296* | *Predicted gene 14296* | 8.527 | 3.67E-46 |
| ENSMUSG00000078867 | *Gm14418* | *Novel KRAB box and zinc finger, C2H2 type domain containing protein* | 8.900 | 4.92E-15 |
| ENSMUSG00000095362 | *Gm14325* | *Predicted gene 14325* | 9.554 | 6.34E-36 |
| ENSMUSG00000078877 | *Gm14295* | *Predicted gene 14295* | 10.562 | 8.04E-22 |
| ENSMUSG00000078862 | *Gm14326* | *Predicted gene 14326* | 10.943 | 1.69E-23 |

**Table S3. Primers for genotyping by PCR.** The primer sequence, amplicon size and reference are listed. bp = base pairs; For = forward; Rev = reverse; neo = neomycin.

| Primer | Sequence 5`→ 3` | Amplicon Size (bp) | Reference |
|---|---|---|---|
| 3`neo* | GATTCGCAGCGCATCGCCTT | WT: ** & *** 222<br>KO: * & *** 636 | This work |
| *Bcan* ForWT** | TATTAAGGAGGAGCGCCGTG | | |
| *Bcan* Rev*** | TCACCCCCTATCATGGGGAA | | |
| *Ncan* ForWT | TCTTGGGGATGCCACGATTC | WT: 331 | This work |
| *Ncan* RevWT | GGGAAACTCCACTGCTGGTTA | | |
| *Ncan* ForKO | GAATCCCCACTCTGCCCTTT | KO: 1,115 | |
| *Ncan* RevKO | ACCTGAGTCAGAGGTAGGGG | | |
| *Tnc* For* | CTGCCAGGCATCTTTCTAGC | WT: * & *** 435<br>KO: ** & *** 342 | [95] |
| *Tnc* neo For** | CTGCTCTTTACTGAAGGCTC | | |
| *Tnc* Rev*** | TTCTGCAGGTTGGAGGCAAC | | |
| *Tnr* For* | AACTCCATGCTGGCTACCAC | WT: * & *** 420<br>KO: * & ** 429 | [82] |
| *Tnr* neo Rev** | ACCGCTTCCTCGTGCTT | | |
| *Tnr* Rev*** | TTTTGGGGAGGTTGATCTTG | | |

**Table S4. Primers for RT-PCR and the generation of *in-situ* riboprobes.** The primer sequence, amplicon size, GenBank accession number and reference are listed. bp = base pairs; ISH = *in-situ* hybridization.

| Primer | Sequence 5`→ 3` | Amplicon size (bp) | GenBank accession number; reference |
|---|---|---|---|
| *Bcan*_ISH_forward | TCAAGTGGACCTTCCTGTCC | 1,093 | NM_007529; This work |
| *Bcan*_ISH_reverse | GACTCGGTAGGTGGTGCAAT | | |
| *Ncan*_ISH_forward | TGCCACGCTCTACACTTGTC | 1,062 | NM_007789; This work |
| *Ncan*_ISH_reverse | GGCTGCATAAGCAGTCATCA | | |
| *Tnc*_ISH_foward | CCATGGGTTCTCCGAAGGAAA | 1,211 | NM_011607; [95] |
| *Tnc*_ISH_reverse | GGATACAGTGGAACAGCAGGT GACGTCACC | | |
| *Tnr*_ISH_foward | GATAGCCCCATGGAT | 1,215 | NM_022312; [95] |
| *Tnr*_ISH_reverse | GAATTTCAAGG | | |
